# Non-uniform Crowding Enhances Transport: Relevance to Biological Environments

**DOI:** 10.1101/593855

**Authors:** Matthew Collins, Farzad Mohajerani, Subhadip Ghosh, Rajarshi Guha, Tae-Hee Lee, Peter J. Butler, Ayusman Sen, Darrell Velegol

## Abstract

The cellular cytoplasm is crowded with macromolecules and other species that occupy up to 40% of the available volume. Previous studies have reported that for high crowder molecule concentrations, colloidal tracer particles have a dampened diffusion due to the higher solution viscosity. However, these studies employed uniform distributions of crowder molecules. We report a scenario, previously unexplored experimentally, of increased tracer transport driven by a non-uniform concentration of crowder macromolecules. In gradients of polymeric crowder, tracer particles undergo transport several times higher than that of their bulk diffusion rate. The direction of the transport is toward regions of lower crowder concentration. Mechanistically, hard-sphere interactions and the resulting volume exclusion between the tracer and crowder increases the effective diffusion by inducing a convective motion of tracers. Strikingly, soft deformable particles show even greater enhancement in transport in crowder gradients compared to similarly sized hard particles. We propose a model that supports the data and quantifies a “diffusiophoretic buoyancy force” when a tracer is in a gradient of crowder concentration. Overall, this demonstration of enhanced transport in non-uniform distributions of crowder is anticipated to clarify aspects of multi-component intracellular transport.

## Introduction

The purpose of this article is to show a counter-intuitive experimental result: We find *higher* transport rates of tracer particles in concentrated solutions of a second species (crowder), when the crowder has a gradient. Further, we suggest a model to describe the physics. Transport of a tracer species through a solution having a gradient of crowder species is a problem of widespread importance. The cellular cytoplasm is highly concentrated with crowing agents such as bio-macromolecules that occupy up to 40% of the available volume, causing the cytosol to be a highly viscous environment^1,2^. The macromolecular crowding has been shown to influence several routine cell activities such as stabilizing enzymatic activity, causing protein binding and aggregation, promoting RNA and protein folding, and stimulating protein-DNA interactions in DNA replication and chromatin organization^3,4,5,6,7^.

Numerous studies in the past few decades have shown that the diffusion of a dilute tracer species always decreases in solutions having a high concentration of crowder molecules. This is often called dampened or hindered diffusion, mainly because crowding increases the viscosity^8,9,10,11^. There are some recent studies that have shown that tracer diffusion coefficients can be close to dilute bulk values, but only at small length scale and when the tracer is much smaller than the crowder so that the tracer sees essentially bulk solution^12,13,14,15,16^. Over longer distances however, diffusion has always been shown to be dampened. These prior experimental studies were done in solutions having a *uniform concentration* of the crowding agent.

Recently it has been shown experimentally that crowding in the cell is heterogeneous with *non-uniform distributions* of macromolecules, especially in eukaryotic cells with inner compartments^17–23^. And so we ask this important question: How is the transport of a dilute tracer species affected in crowded systems that resemble the intracellular cytoplasm, where the crowder species has a *concentration gradient*? To our knowledge, there are only two modeling-based papers in the literature that have looked into the tracer motion under crowder gradient^24,25^. However, there are no reported experimental results for tracer transport in a gradient of crowder molecules, and this has prevented the assessment and applicability of these two papers for non-equilibrium crowded conditions.

In this paper we experimentally study the transport of hard and soft-matter probe tracers that reside in either uniform or non-uniform concentrations of crowder macromolecules. This was enabled by our development of a new measurement method for these systems. Our experiments involved three components: tracer, crowder, and buffer solution. We compare the tracer’s transport in a gradient of crowder to the transport in uniformly-crowded media. Benefiting from a microfluidic system, we investigated the transient dynamics of tracer transport under these gradient conditions. We further explored key factors affecting the tracer transport, including 1) size of the tracers, 2) crowder concentration, and 3) crowder molecular weight. We further explore a key factor for biological cells: 4) tracer deformability using hard structures (i.e., polymer microspheres) and soft structures (e.g., proteins, polymer macromolecules, and protocells, mimicking cellular organelles).

Our experimental results contradict parts of the aforementioned models as will be explained later in this article ^24,25^. Therefore, we developed our own model, applying the well-known theory of non-electrolyte diffusiophoresis. We find that this model explains our experimental results much better, accounting appropriately for viscosity. In developing our model, we introduce the concept of “diffusiophoretic buoyancy”, which provides a quantitative and intuitive way to think about the tracer transport in a gradient of crowder. We anticipate that our experimental and modeling results will have a significant impact on understanding transport in crowded biological cells.

## Results

Our results encompass several key outcomes: 1) Detection of enhanced tracer transport in a gradient of crowder, 2) effect of concentration and particle sizes of both tracer and crowder, 3) comparisons of hard and soft tracers, 4) new model for tracer transport in a gradient of crowder, and 5) enhanced diffusion when the tracer and crowder have the same size.

### Measuring the Transport Distance of Tracers

We experimentally studied the transport of tracer colloids (i.e. soft macromolecules and hard particles) in crowded environments of macromolecules under two different conditions: uniform and non-uniform distributions of crowder. In the uniform crowder condition, we measured the diffusion coefficient of the tracers with Fluorescent Correlation Spectroscopy (FCS) which is a method that has been previously established in studying transport in crowded environments^10,11^. In addition, we utilized microfluidics to establish stable gradients of crowding macromolecules as a new technique to study the tracer transport under non-uniform crowding conditions.

The experimental setup involved a microfluidic device that enabled us to have stable gradients of crowder. We used a three inlet-one outlet polydimethylsiloxane (PDMS) microfluidic channel with dimensions of 4 cm length × 360 μm width × 50 μm height (Figure 1A) with solutions pumped in through each inlet at 50 μl/hr (See Methods). The solution of macromolecular crowder in buffer enters through the middle inlet while only buffer (1 mM KCl) enters through the side inlets. At this flow rate, the average residence time of the solutions within the main channel is 17.4 s from the mixing point to the end. This setup produces a symmetric concentration gradient of crowder macromolecules across the main channel with the most crowding in the middle of the channel and least crowding at the side walls. Along with the crowder, fluorescently-tagged probe tracers (either tracer macromolecules or polystyrene beads) were also added to the solution entering the middle inlet. We measured the concentration profile of the tracer particles using confocal microscopy near the end of the channel (39 mm) at the mid-depth (~20 − 25 μm up from the bottom) (Figure 1A). For control experiments, only the tracers were added to the mid-inlet solution and diffuse toward the sides as they travel with the fluid down the channel. The normalized intensity profile of tracers across the channel near the end is shown in Figure 1B for the specific case of 60 nm cPSL tracer particles (carboxylate-modified polystyrene) without and with poly(ethylene oxide) 1000 kDa at 0.3 wt% (weight percentage) as the crowding species in the mid-inlet solution. Note that the normalization step is done to make comparisons possible between different samples. Normalization is done by setting the minimum and maximum intensity to zero and one, respectively. The corresponding non-normalized curves along with the conservation of mass balance calculation is provided in SI. There are two differences between the two curves: (i) The slope of the tracer profile at the half-maximum intensity, the inverse value of which gives the lateral transport of the 60 nm tracer particles. This value of lateral transport distance is the main parameter indicating the crowder’s effect on the tracer’s movement. (ii) a broadening of the experimental curve toward the sides (shown with brown arrows). This broadening is due to a hydrodynamic effect resulting from the higher relative viscosity of the polymer solution in the mid-inlet compared to buffer. When this viscous fluid is flowed from the mid-inlet while the buffer enters from the side-inlets, the middle flow pushes the buffer to the sides hydrodynamically. This effect takes place immediately after the mixing point in the main channel and is not indicative of the type of molecular interaction we discuss here (see SI). Therefore, this broadening does not influence the experimentally measured values of transport distance of tracers in crowder gradients.

**Figure 1.**
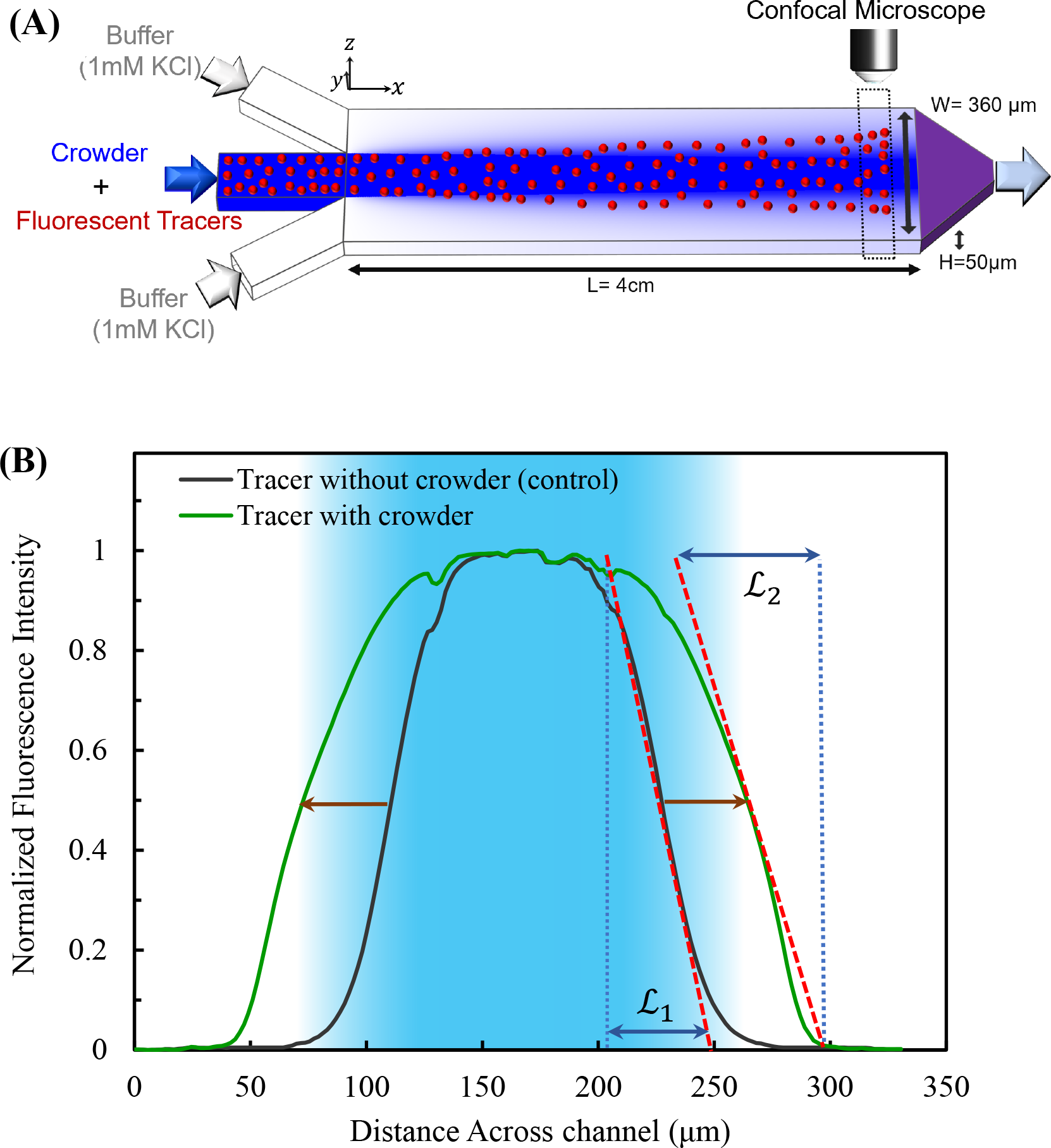
Experimental setup and the resulting profile used for measuring the transport of tracers. **(A)** A three-inlet one-outlet polydimethylsiloxane (PDMS) microfluidic device with dimensions (L × W × H) of 4 cm × 360 μm × 50 μm was used to measure the transport of tracers. Fluorescently-tagged tracer species, polystyrene beads or macromolecular tracers, along with macromolecular crowder is pumped through the middle inlet while buffer is pumped through the side inlets. This setup generates a lateral concentration gradient of crowding species across the channel with more crowder in the middle of the channel and less crowder near the side walls. This crowder gradient affects the motion of the tracer species, mimicking the heterogeneously crowded cytosol environment. In all the experiments, the flow rate entering each inlet is maintained at 50 μL/hr using a syringe pump. To obtain the intensity profile of the tracers, confocal scanning microscopy was performed near the end of the channel (~39 mm from the inlet = ~ 17 seconds of interaction time). All the solutions are in 1 mM KCl. **(B)** The normalized fluorescent intensity profile of 60 nm cPSL tracers is obtained at the end of the channel in the absence and presence of PEO-1000kDa crowder in the mid-inlet. After 17.4 seconds of residence time within the channel, the particles in the mid-flow diffuse out with a transport distance of 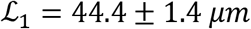 (black curve) corresponding to its diffusion coefficient of 9.0 ± 0.6 *μm*^2^/*s*. However, in the presence of the crowder gradient (blue background shade) of 0.3 wt% PEO-1000 kDa, the tracers show a larger transport distance, 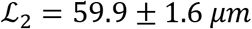 (green curve) despite their dampened brownian diffusion. The resulting enhancement in the transport distance is 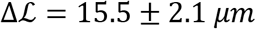. The brown arrows show broadening due to a hydrodynamic effect which does not impact the lateral transport distance. The graphs are obtained by averaging three videos each with 633 frames over 5 minutes.

### Tracer Transport Increases in a Non-Uniform Crowded Environment

We observe a higher transport rate in a crowded environment where we might normally expect to see a lower rate due to enhanced viscosity effects. In the absence of the macromolecular crowder in the mid-flow, the probe tracers move along a distance which corresponds to the diffusive length scale. This distance can be obtained by taking the inverse of the fluorescence curve’s slope at 0.5 intensity (Figure 1B). We define this length as the *transport distance* 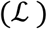 throughout this paper (See Methods for video analysis). In figure 1B, the initial transport distance is 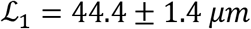. Based on this transport distance, we can calculate the diffusion coefficient of the tracer as follows (derivation in SI):

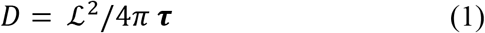

where *D* is the diffusion coefficient, 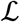 is the transport distance and **τ** is the average residence time in the microfluidic channel which is equal to 17.4 s for our microfluidic experiments. For this specific case, 60 nm cPSL tracers in the absence of any crowder, the lateral transport distance is 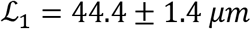 which would give the diffusion coefficient of 9.0 ± 0.6 μm^2^/s. This diffusion coefficient from microfluidics is close to the diffusion coefficient we obtained by FCS 8.2 ± 0.5 μm^2^/s corresponding to a 60 ± 4 nm particle. In the presence of the crowding polymer in the mid-flow, 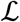 increases to 59.9 ± 1.6 μm. Using the same calculation, we can obtain an apparent diffusion coefficient of 16.4 ± 0.9 μm^2^/s in the presence of the crowder in the mid-flow, about twice that of the base diffusion coefficient. In contrast to this diffusion enhancement in a gradient, in the presence the same crowder under uniform 0.3 wt% concentration, the tracer diffusion measured by FCS shows a significant decrease to 1.1 ± 0.1 *μm*^2^/*s* (>85%). Considering the diffusion enhancement in non-uniform and diffusion drop in uniform distributions of the crowder, the striking question regarding the source of this difference arises. We investigated the underlying mechanism behind the enhanced transport observed in non-uniform crowding. As previously stated, we hypothesize that the hard-sphere repulsive interaction in the form of the excluded volume effect may lead to the enhanced transport of the tracers.

### Factors Affecting the Tracer Transport in a Non-Uniform Crowded Environment

We showed that the transport distance of the 60 nm hard sphere tracers increase upon addition of PEO-1000kDa to the mid-inlet solution as the macromolecular crowder. We can express this increase in terms of *enhancement in transport distance*, 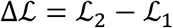, where 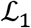 and 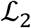 are the transport distances in the absence and presence of crowder, respectively. In other words, 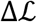 is the measure of how much the crowders “push” the tracers toward the sides where they are less populated. The enhancement in the transport distance for the 60 nm tracers is 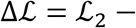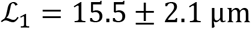. Here, we have varied the size of the tracers from 15-540 nm to investigate the effect of tracer size on its transport distance. Figure 2 shows the enhancement in the transport distance, 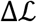, for tracer beads with various sizes. As can be seen, 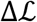 is close to zero for 15 nm tracers and showing that their motion is not affected by the presence of the PEO crowding molecules. The negligible transport enhancement of 15 nm tracers can be attributed to their size being significantly smaller than the size of the crowder (PEO-1000kDa diameter is measured to be 96.9 ± 9.1 nm by DLS). For 35 nm diameter PS tracer particles, the presence of the PEO crowder starts to affect the tracer transport, enhancing its transport by 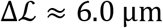. This effect of enhanced transport further increases for larger tracers, ~60 *nm*. However, particles larger than ~60 *nm* experience similar enhancement in transport distance. The similar values of 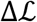 for large tracers, especially larger than the crowder, indicates that the macromolecular crowder in the middle pushes the tracer toward the sides to the same extent if the tracers are larger than the crowders, independent of tracer diameter. Therefore, if the size of the hard tracer exceeds the average interstitial size between the macromolecular blobs of the crowded medium, the tracers “feel” a pressure from the crowding polymers in the middle and as a result, they are dislodged from the microenvironment at an increased rate due to repulsion by the crowders. Note that this effect is independent of the charge of the colloidal tracers (see SI), but depends on the interstitial size dictated by correlation length of the crowder ^26^. We also looked at the case with a different initial crowder distribution where the crowder solution is being flown from the side-inlets while tracers are only present in the middle. We saw that the tracers are also pushed away from the crowder and toward the middle by an amount that is equal and opposite compared to the case of crowder in the middle (see SI). Similar focusing behavior has been observed when the initial concentration of tracers is uniform as well (See SI).

**Figure 2.**
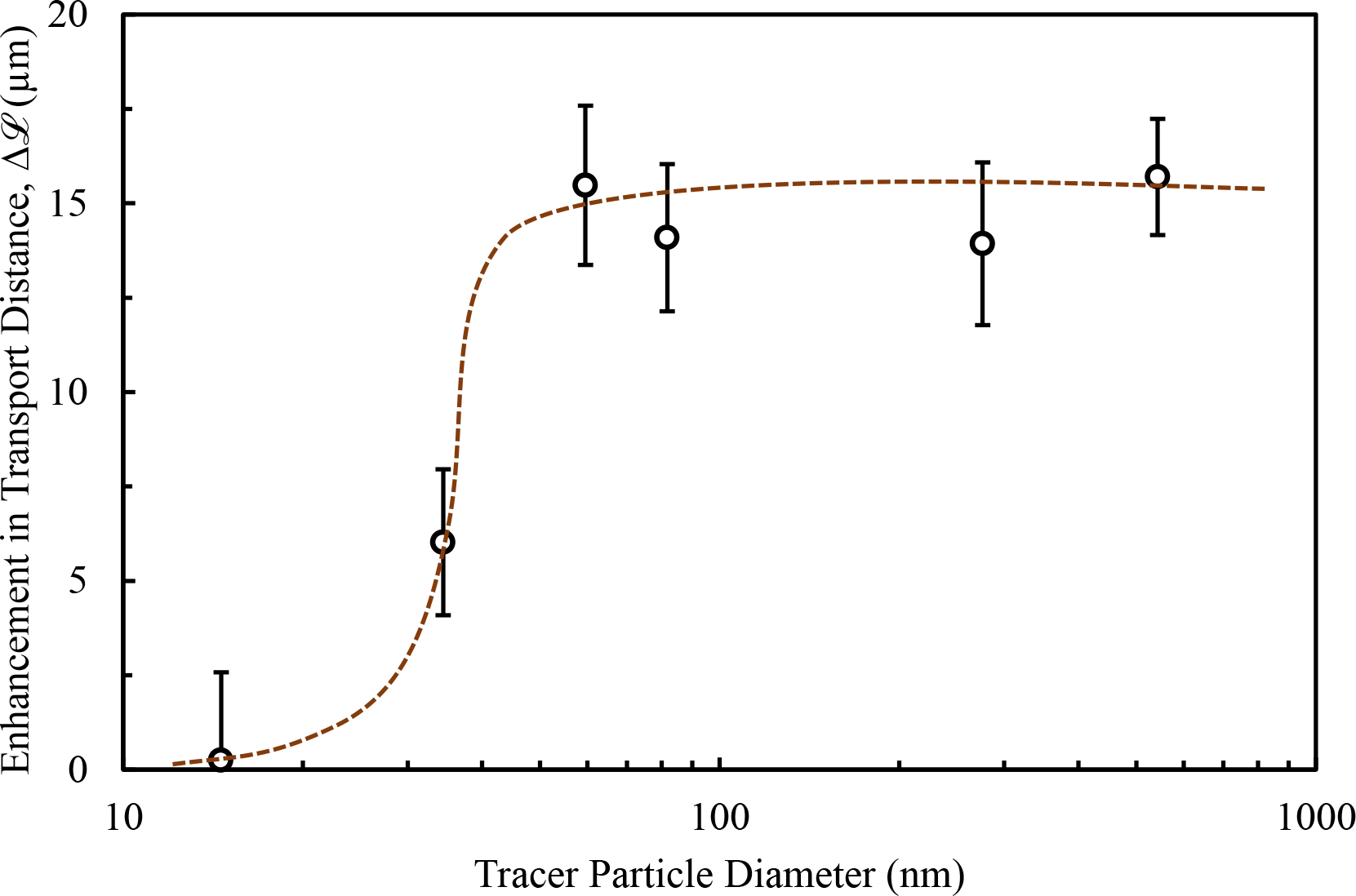
Effect of the diameter of hard tracer particles on the enhancement in their transport distance with the presence of the 0.3wt% PEO-1000kDa (diam. size ≈ 97 nm) crowder gradient. Enhancement in transport distance, 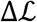, is calculated from the difference between the tracer transport distance in the presence of the crowder, 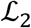, minus that of control, 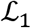. The transport of smaller tracers is not significantly affected by the presence of the macromolecular crowder. For larger tracers however (> 60 nm), values of 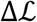 are almost similar, indicating that the tracers “feel” similar pressure from the crowding polymers in the middle when the size of the tracer itself is comparable with the size of the crowders indicating of hard sphere repulsions. All the tracers are polystyrene beads except 15 nm silica particles (See tracer sizes in SI). The 90% CI error bars are obtained by averaging the slope from three profiles each obtained from a video with 633 frames over 5 minute.

The next variable we probed was how the size of the macromolecular crowder impacts the transport of the tracer. At similar weight fraction of PEO polymer crowder of 0.3wt% in the mid-inlet solution, we varied the molecular weight from 20 kDa (diam. size = 12.7 ± 1.6 nm) to 2000 kDa (diam. size = 116.0 ± 8.9 nm) with the same colloidal tracer: 82 nm amine-modified PS particles (aPSL). The enhancement in the tracer transport distance, 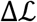, is depicted in Figure 3A. As can be seen, the presence of the PEO crowder with sizes of 20 kDa and 200 kDa has almost no impact on the tracers’ transport, giving 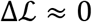. However, higher molecular weight PEO crowders with larger sizes push the tracers toward the sides more. Hence, the enhancement in the transport distance of the tracers, 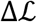, increases monotonically with the size/molecular weight of the polymer. For 2000 kDa PEO (diam. size= 116 nm), we observe a very different pattern of the aPSL tracer profile than the other cases as depicted in Figure 3B. The green curve consists of a narrow and a wide region: (*α*) The diffusive region, where there is almost no crowding polymer molecules; and (*β*) ultra-excluded volume region dominated by crowding effect, where we have the highest molecular crowding that pushes the tracer particles away from the middle toward the sides. Therefore, we can observe diffusion-dominated as well as crowding-effect dominated regions in the same curve. The reason that this specific case of 2000 kDa PEO shows a different pattern is because the tracer particle is smaller than the crowder and has a higher diffusion coefficient. Although both tracer and crowder start diffusing out from the middle region, the tracers surpass the crowders to form the diffusion-dominated region *α* where there is almost no crowder present. The rest of the tracer particles are being pushed out strongly by the large PEO-2000kDa crowders, forming the wider region *β*.

**Figure 3.**
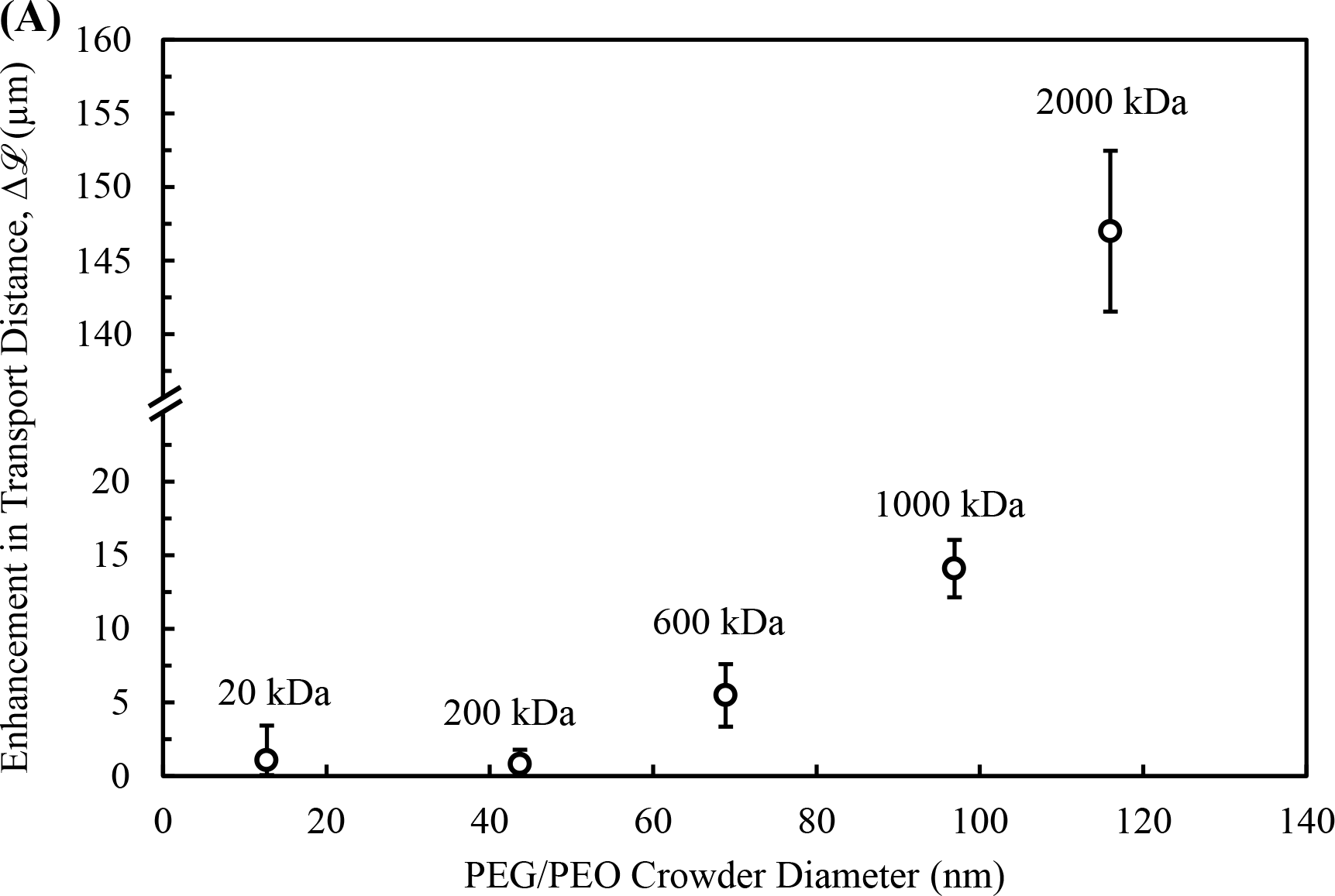
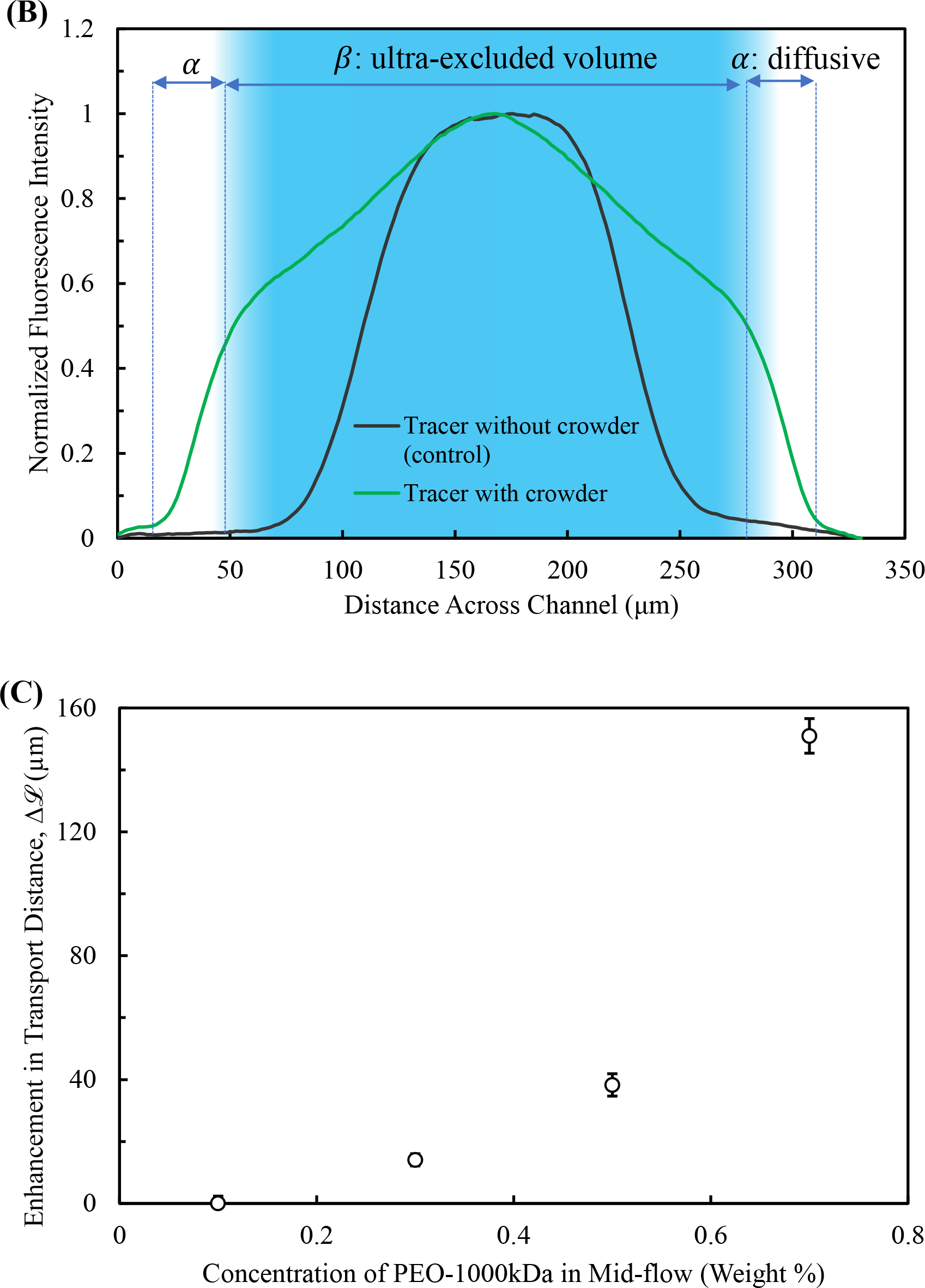
The effect of varying crowder size and concentration on the tracers’ motion. (A) Effects of crowder 0.3 wt % PEO size on the transport distance of tracer 82 nm aPSL particles. As the size of crowder PEO increases, the enhancement in transport distance significantly increases. (B) The tracer profiles observed from 2000 kDa PEO in Figure 3A with (*α*) the diffusion-dominated region, where there is almost no crowding polymer; and (*β*) the crowding-effect dominated region where we have the highest molecular crowding that pushes the tracer particles away from the middle toward the sides. The shadowed area is where the polymer molecules is expected to be present (C) Shows the effect of varying concentration of crowder PEO-1000kDa concentration on transport distance of tracer 82 nm aPSL particles. As the concentration of crowder 1000 kDa PEO increases, the transport distance is observed to increase exponentially. The 90% CI error bars are obtained by averaging the slope from three profiles each obtained from a video with 633 frames over 5 minute.

In addition to the size of tracer or crowder, another factor affecting the tracer transport is the crowder concentration. Higher concentrations of macromolecular crowder are expected to push the tracer particles more, leading to larger transport of the tracers to the side. Figure 3C is the enhancement in the transport distance of 82 nm tracers in the presence of PEO-1000 kDa (diam. size ~97 nm) in the mid-inlet solution by varying the polymer concentration up to 0.7 wt%. As can be seen, higher concentration of crowder leads to more tracer transport toward the sides. Tracer transport is exceptionally high at 0.7 wt% of polymer due to the very high crowder concentration. This concentration is more than 4 times of the overlapping concentration, C* = 0.14 wt%, the concentration at which the polymer macromolecules are densely packed. Note that at 0.7 wt%, the tracer profile forms a pattern as shown in Figure 3B. These similar profiles show that both large size and high concentration of crowder, especially exceeding the overlapping concentration of the macromolecule, dramatically increases the observed transport enhancement due to crowding-effect.

### Comparing the Enhanced Transport of Hard vs. Soft Matter Tracers

Until now, all the experiments have involved hard colloidal tracers in soft-matter macromolecular crowders (PEG/PEO polymers). To mimic further the biological environment where all the species are soft-matter (i.e. proteins, organelles and cellular compartments), we explored soft-matter tracers and compared them with the previous results for hard colloids.

As can be seen in Figure 4A, the soft tracers (urease, dextran, vesicles) experience higher 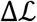 compared to the hard tracer particles in all cases when comparing similar sizes. Both 500 kDa and 2000 kDa dextran show a 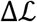 of ~27 μm while the vesicles show a 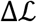 of ~35 μm. The respective PSL hard matter all show a 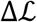 of ~15 μm. Although the soft tracers show larger enhancement in transport, the “*transition size*” is similar, the size where transport enhancement dramatically increases (Figure 4A). This transition size is ~35 nm for both types of colloids, presumably the same as the pore size of the PEO crowder solution. According to the principle of volume exclusion, particles larger than this characteristic interstitial size of the crowder milieu would be immediately expelled, which corroborates our experimental observations.

**Figure 4.**
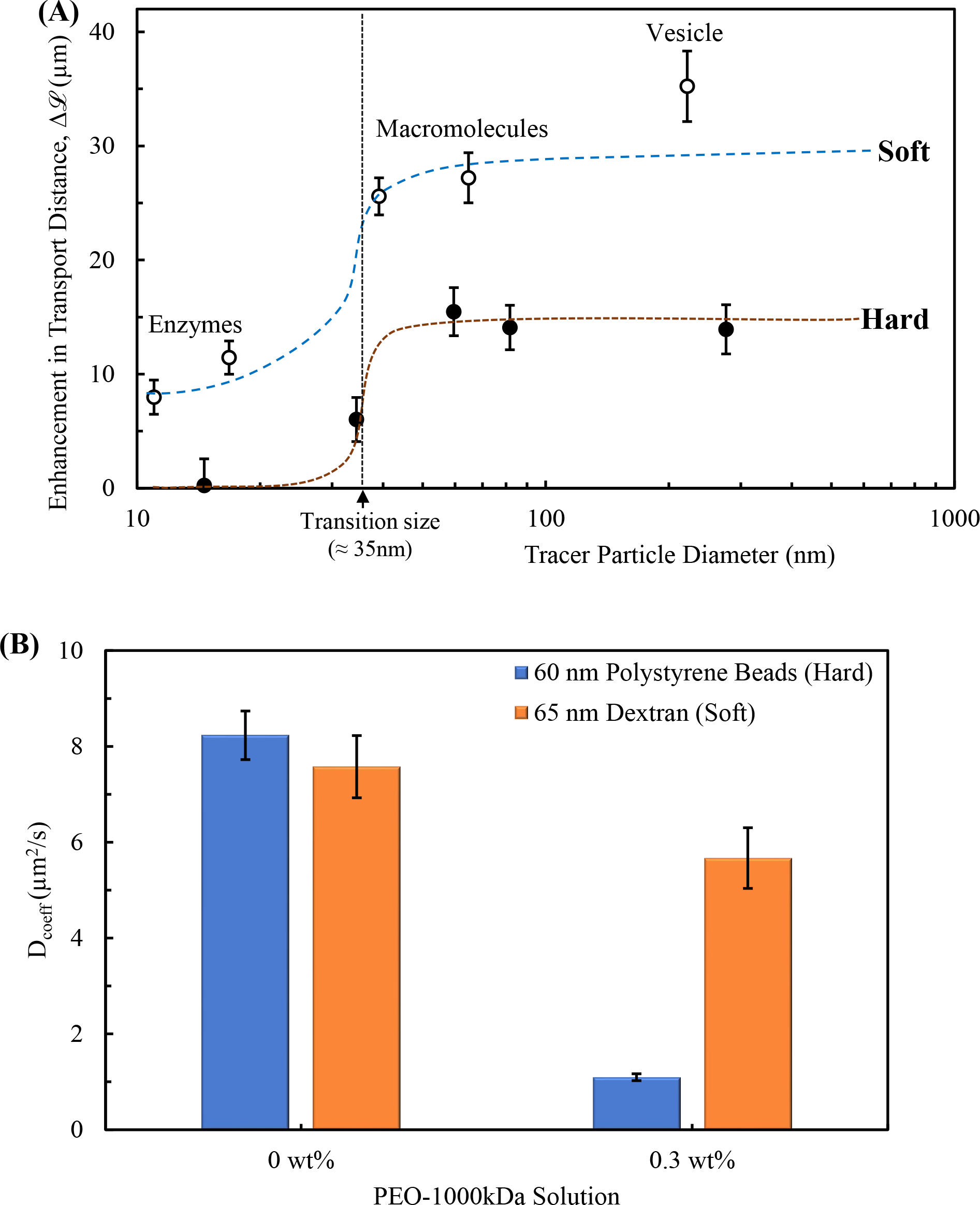
Comparing the transport distance of similarly sized hard and soft matter. (A) Enhancement in the transport distance of similarly sized hard and soft matter, all in a similar gradient of crowder: 0.3 wt% 1000 kDa PEO. In general, soft tracers shows a higher transport distance compared to hard tracers in non-uniform crowding. However, the “*transition size”* is similar for both types, the size where transport enhancement dramatically increases. (B) FCS data of 2000 kDa Dextran (soft tracer) and 60 nm aPSL (hard tracer) with and without uniform crowder 0.3 wt% 1000 kDa PEO. The diffusion of hard tracers drops significantly while the diffusion of soft tracers is only slightly decreased in uniform crowding due to compression of the radius of gyration. Note that the error bars are based on 90% CI for at least 8 FCS readings.

In order to find out the reason for soft tracers showing greater enhanced transport compared to hard tracers, we have looked at the diffusion of one set of soft vs. hard tracers in equilibrium crowder by FCS measurements (see Methods). Figure 4B shows FCS data for 2000 kDa dextran and 60 nm PSL particles in the presence and absence of 0.3 wt% of 1000 kDa PEO crowder. The diffusion of both species is almost equal in buffer due to their similar size. However, in the crowded environment, polystyrene particles experience a significant drop in diffusion, more than 85%, while 65 nm 2000 kDa dextran diffusion decreases minimally. The significant drop in diffusion of the hard particles are due to the increased microviscosity of the PEO solution. It is to be noted here that if the tracer particle radius remains smaller than the radius of gyration of the crowder, the former would only feel microviscosity of the solution. The soft matter tracers are more deformable compared to the rigid hard spheres and can squeeze into the interstitial spaces of the crowder to retain near normal diffusion while the hard matter cannot undergo compression and experiences the macroviscosity of the medium^12,27^. In a non-uniform crowded environment, both the hard and soft matter experience the same force from the crowded region but the soft matter is deformable, being able to move away from the crowder faster compared to the hard matter similar to their diffusion trend (see SI). The vesicles show even more enhanced transport compared to the other macromolecular soft matter as the hollow vesicles are expected to be more deformable compared to the polymer macromolecules. So, deformability of the tracer species is very important in the level of their enhanced transport in non-uniformly-crowded environments, determining how fast the tracers can “escape” from the more crowded regions.

### Proposed Model for Transport Under Non-Uniform Crowding

We experimentally observed that a relatively large tracer particle is pushed out of a crowded region in the presence of a gradient of macromolecular crowder (Figure 4A). A few models have been proposed to explain such behavior. Schurr et al. proposed a model based on the occupied and inaccessible volume by the crowder and predict a linear dependency of drift tracer velocity to its size at larger sizes, while we experimentally observed a plateau^24^. Using the mean-field approach, Smith et al. derived a transport equation under heterogeneous condition for the transport of both tracers and crowders^25^. The proposed equation however, suggests that the tracer drift velocity is inversely proportional to the tracer size at smaller size ranges, therefore transport dramatically increases at small sizes. We experimentally observed the opposite, as the smaller tracer showed much lower transport distance compared to larger tracers (Figure 2). A theory for collective diffusion has been also developed; however, these models have been developed for two species (i.e. one species in liquid), where the key changes happen at higher concentrations of tracer particles ^28,29^. Our systems have three components and the proposed theory is based on a different physics which can be applied to multi-component systems that is relevant to biological cells.

An alternative perspective can be taken, considering the movement of the tracers as a diffusiophoretic motion in a gradient of hard sphere crowders. A rigorous theory has been developed by Anderson, Prieve, and co-workers for the motion of a colloidal particle in the gradient of interacting solutes^30–32^. This theory can be applied to our system as well, considering the tracers as the colloidal particles and crowders as the interacting solutes. Under that condition we can use a variation of this theory where the two species interact by “hard-sphere repulsion”. The hard-sphere repulsion of the crowders induces a fluid slip velocity at the surface of the tracer which moves the tracers away from the high concentration of the crowding species. The escaping/drift velocity of the tracer particles from the high concentration of crowder, U_escaping_, can be obtained from the following equation:

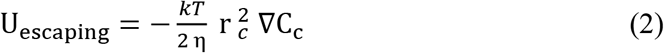

Where *k*T is the thermal energy; and *r*_*c*_ and ΔC_c_ is the hydrodynamic radius and concentration gradient of the crowder species. *η* is the viscosity of the solution, which is often highly dependent on the concentration of the crowder as the crowder increases viscosity. Here, the concentration of the tracer species is kept so low to the extent that it would not impact viscosity or other solution properties.

The assumption behind the diffusiophoretic theory is that the tracer size should be much larger than the radius of the crowder, *r*_*tr*_ ≫ *r*_*c*_. Under that condition, the drift velocity of the tracer species is fully independent of the size of the tracer and depends only on the size and concentration gradient of the host crowding species. This statement agrees with the experimental data. According to figures 2 and 4A, the enhancement in the tracer transport is relatively similar for large tracers, larger than the transition size which is ~35 *nm*. For smaller tracers however, experimental results show that the tracer motion goes to zero as the size of it decreases. Unfortunately, the theory has not yet been extended for the small tracers since the assumption of the theory does not hold for that size range.

In addition to accounting for tracer and crowder size, the theory can also link the lower diffusion dampening of soft colloids to their higher enhanced transport compared to hard spheres, i.e. relation between Figure 4A and 4B. As discussed earlier, unlike hard spheres, deformable soft colloids can squeeze into the interstitial spaces in the crowded medium to retain near normal diffusion. So, it is reasonable to state that soft colloids experience a lower micro-viscosity in a crowded environment compared to the higher macro-viscosity that the hard spheres experience^12,27^. Considering the lower viscosity (*η*) that the soft colloids experience when applying equation 2, they would have a higher escaping velocity compared to the hard spheres in a similar condition. In brief, the schematic in Figure 5 summarizes the observed phenomenon under non-uniformly distributed crowders, Small sized tracers do not show any enhanced transport while both large hard and soft colloids escape away from the high crowder concentration at a specific escaping velocity, which is higher for soft colloids compared to hard spheres. The higher escaping velocity leads to larger enhancement in transport of the soft species.

**Figure 5.**
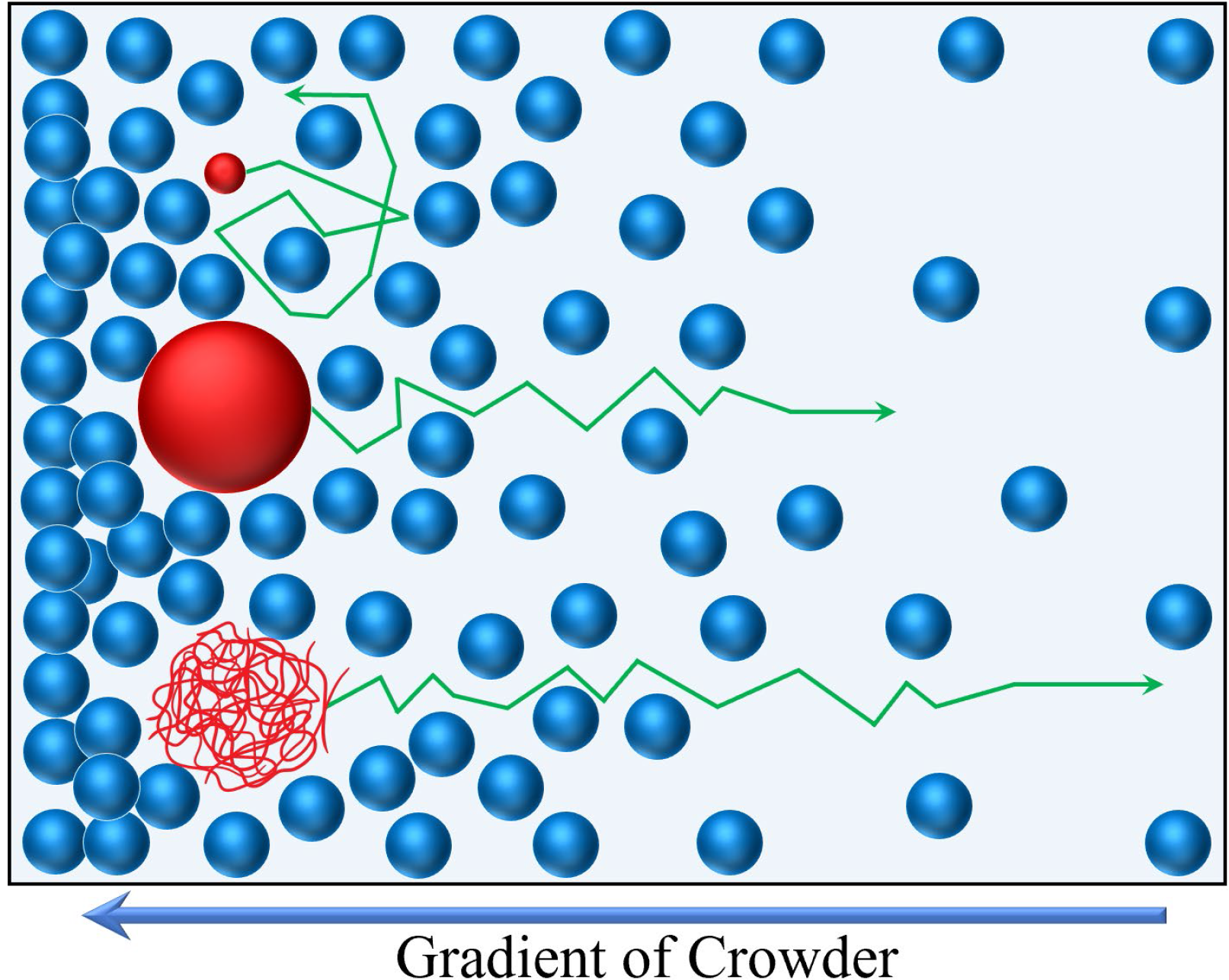
Schematic of the tracer motion under non-uniform crowded environment: The motion of small size tracers tends to remain unaffected under non-uniformly distributed crowders while both large hard and soft colloids escape away from higher crowder concentration at a specific velocity, termed as “escaping/drift velocity”. This velocity is higher for soft colloids compared to hard spheres and as a result, soft colloids move away faster.

### Self-diffusion increases with concentration

In this section, we focus on a specific case where both tracer and crowder are identical soft-matter macromolecules. This case is especially important to explore the changes in the mode and value of self-diffusion at uniform and non-uniform concentrations. The probe macromolecule in this section is dextran 2000 kDa with a hydrodynamic diameter of ~65 nm. FCS was used to measure the tagged-dextran diffusion in a uniformly crowded environment of untagged dextran. Also, we performed similar microfluidic experiments for transport under non-uniform concentrations with a mixture of tagged dextran and untagged dextran 2000 kDa at various concentrations in the middle inlet (Figure 6A).

**Figure 6.**
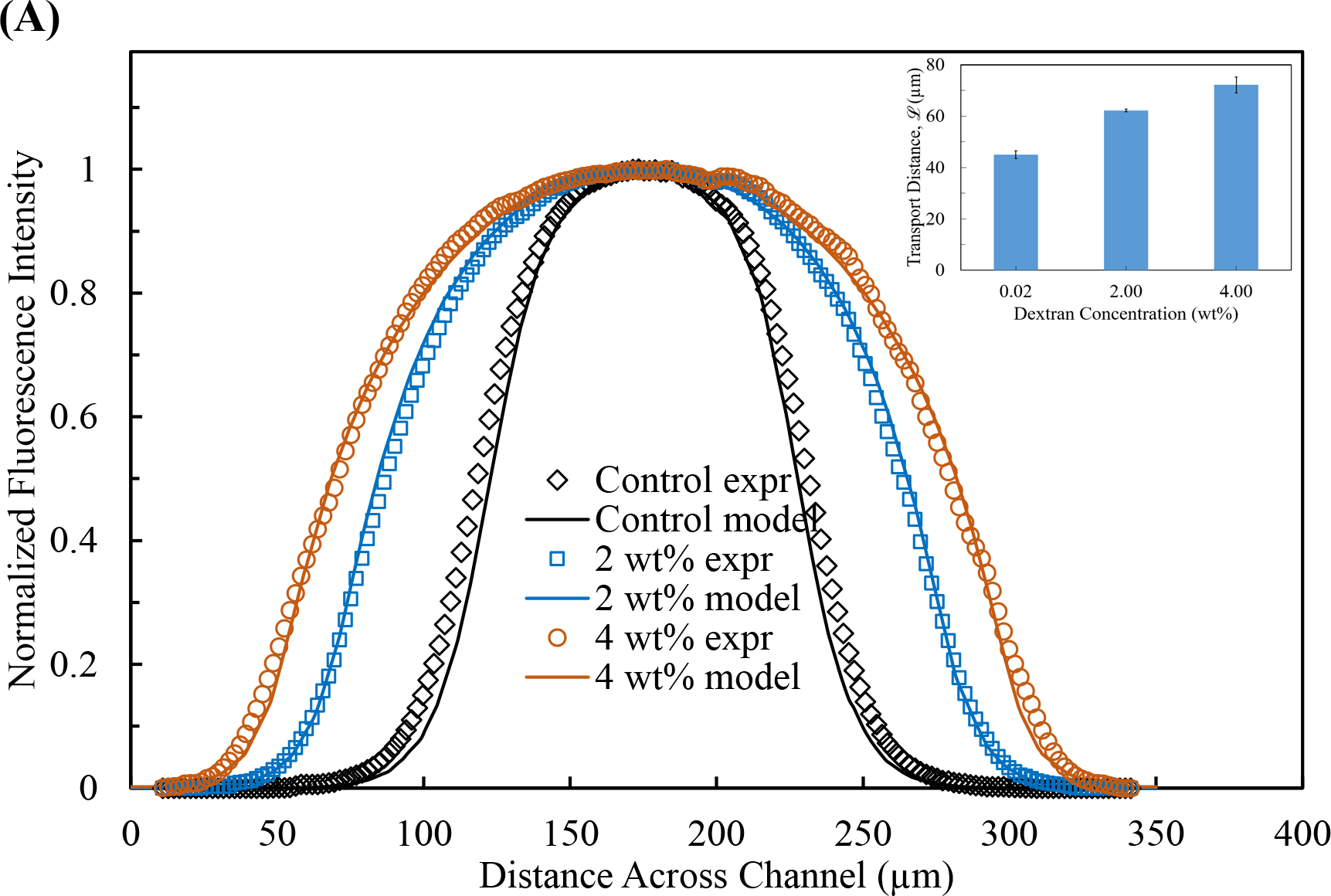
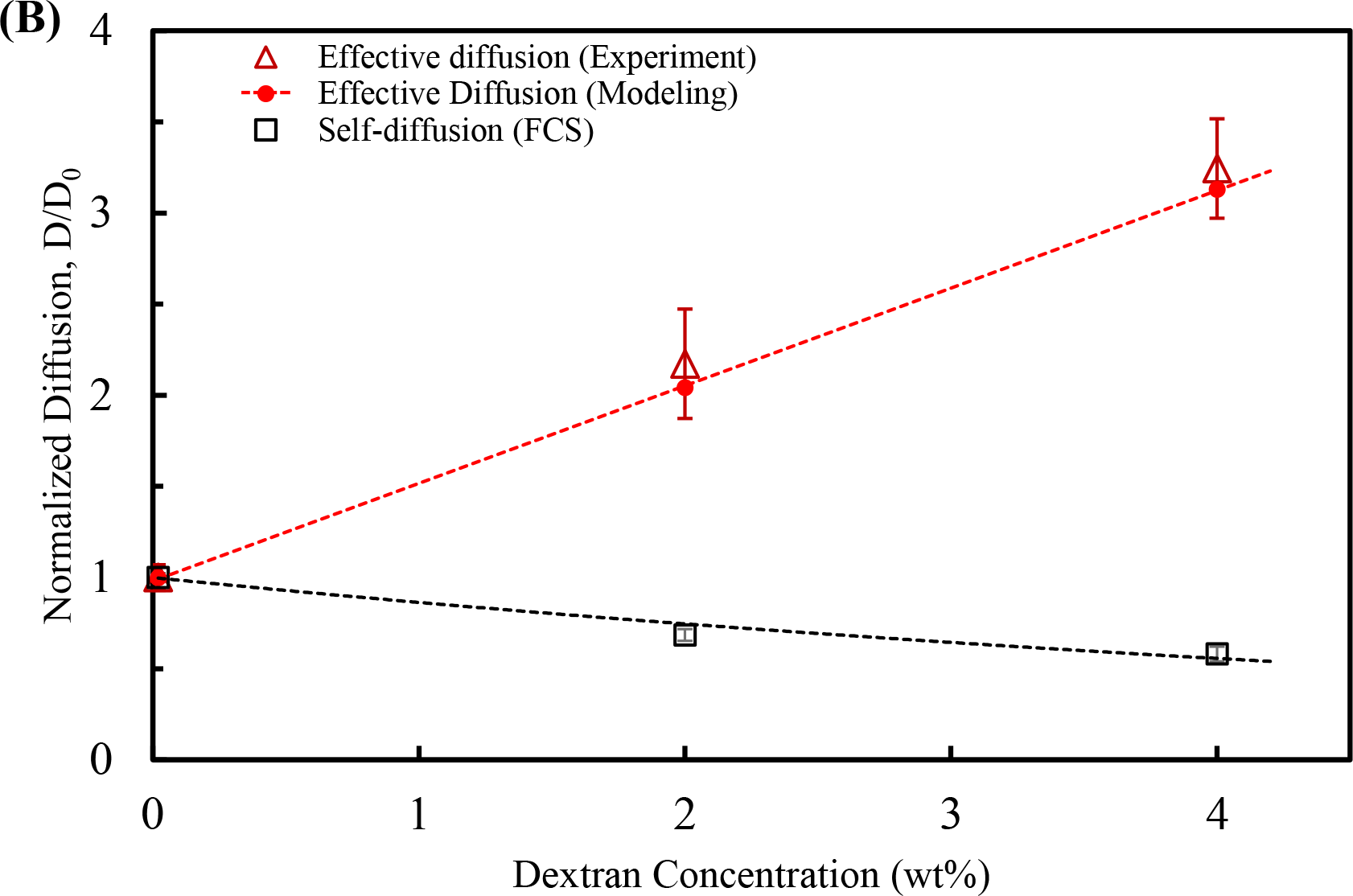
The self- and effective diffusion of 65 nm Dextran tracer. (A) Experimental and modelling curves of dextran intensity profiles at the end of the channel when 0.02 wt% (control), 2 wt% and 4 wt% dextran are in the middle flow. Inset: the corresponding lateral transport distance of the experimental curves at the 0.5 intensity. As can be seen, the transport distance increases with higher dextran concentration in the middle inlet. (B) The self-diffusion of dextran macromolecules drops at higher concentration of themselves while their effective diffusion increases in non-uniform initial concentration. Note that the error bars on self-diffusion values are based on 90% CI for at least 8 FCS readings.

We measured the self-diffusion of the fluorescently labeled dextran at different concentrations of untagged dextran via FCS (black squares in Figure 6B). As expected, a moderate decrease is observed. We fitted the self-diffusion data to the phenomenological functional form below from the literature, to estimate the diffusion of dextran at different concentrations:^33,12,34^

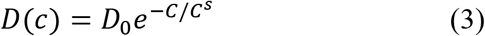

Where D(c) is the self-diffusion coefficient as a function of concentration, and D_0_ is the self-diffusion co-efficient of the dextran macromolecules at infinite dilution which is measured to be *D*_0_ = 7.6 ± 0.6 μm^2^/s (corresponding diameter of ~65nm). C^s^ is a scaling concentration at which the diffusivity drops by 63%. Fitting to the data, we find C^s^ = 6.7 wt% ≡ 33.5 μM (black dash line in figure 6B). We also confirmed the diffusion drop with microfluidic experiments with equal concentration of untagged dextran in all three inlets (see SI)

For the transport under non-uniform concentrations, we performed similar microfluidic experiments mentioned above with a mixture of tagged along with untagged dextran 2000 kDa at various concentrations in the mid-inlet. The normalized intensity profile of the dextran near the end is shown in Figure 6A. The dotted lines are the experimental curves. From low to high concentrations, the slope at the half-maxima decreases which leads to enhancement in the transport distances, 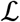. The values are shown in the inset in Figure 6A. The initial transport distance is 43.2 ±1.1 μm for a very dilute concentration of tagged dextran in the middle of the channel, corresponding to a diffusion of 8.5 ± 0.4 μm^2^/s, which is close to the base diffusion measured by FCS. Increasing the dextran concentration in the mid-inlet, the transport distance increases to a peak value of 76.7 ± 1.0 *μm* at 4wt%. Although the diffusion of the dextran is not entirely Fickian in the higher concentration of dextran due to complex crowding effects, we can still calculate an effective diffusion coefficient using equation 1, giving a peak value of 26.9 ± 0.7 μm^2^/s at 4 wt%, which is about three times larger than the diffusion coefficient obtained in the control experiment in the absence of crowder. Therefore, dextran macromolecules show an enhanced diffusive behavior at higher concentration gradients.

In fact, this enhancement is anticipated by accounting for the escaping velocity and the resulting convective transport in crowded medium which can be explained by equation 2. However, here we have only one species as the tracer and crowder species are the same, so we let *C* = *C*_*c*_ = *C*_*tr*_ be the overall concentration of dextran. Therefore, the total flux of the dextran macromolecules can be obtained by summing over the diffusive and convective fluxes as the following:

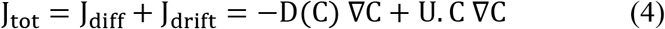

Where D(C) is the self-diffusion coefficient of the macromolecules obtained from equation 3, and C is concentration of the macromolecules. U is the escaping velocity defined in equation 2. Substituting equation 2 and writing *kT*/*η* in terms of diffusion as 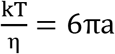 D(C), we can rewrite the equation 4 as the following:

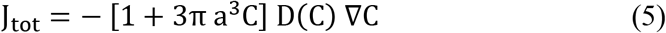

The above expression has a similar form as Fick’s diffusion equation relating flux to the concentration gradient but with a variable coefficient which is dependent on the concentration the diffusing species. By setting the above equation equal to a Fickian form relation, J = D_eff_ ∇C, we can define an effective diffusion coefficient under gradient, D_eff_, as the following:

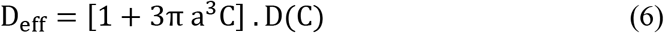

Which is comprised of self-diffusion coefficient *D*(*C*) that decreases as concentration increase (equation 3), as well as another term in the bracket, increasing with concentration. The experimental values of effective diffusion for the profile shown in Figure 6A are plotted in Figure 6B (red curve and triangles). As can be seen, the effective diffusion under dissipating gradients enhances at higher concentrations of dextran, although the self-diffusion (black curve) decreases. This trend is predicted by the above equation and confirmed from microfluidic experiments.

In order to further assess the agreement between the model and experiment, a comprehensive simulation was performed based on the model for escaping velocity in crowded environment. This was done by solving the transport equation 5 for the dextran macromolecules over the three-dimensional geometry of the channel created in COMSOL Multiphysics software (v5.3). This method was utilized in our previous works^35,36^. To obtain the highest accuracy and incorporate the effect of the viscosity, we initially solved the incompressible Navier-Stokes equation to obtain the fluid flow in the channel. Viscosity of the dextran solution was measured using a viscometer (Brookfield DV-II+ Pro) (see SI for viscosity measurements of dextran 1000kDa). Then, the mass transport equation (equation 5) was solved to obtain the concentration profiles (see Methods). The resulting profiles are plotted in Figure 6A (solid lines). As can be seen, very close agreement can be seen between the experimental and modelling data. After obtaining the dextran concentration profiles through simulation, the transport distance and the resulting effective diffusion coefficients are calculated at different concentrations and plotted in Figure 6B (red dotted line). The modelling values are plotted in Figure 6B (red circles). Very good agreement is observed between the experimental data (red triangles) and the simulation (red dotted line).

The effective diffusivity defined in equation 6 has interesting implications. First, it suggests an enhanced-diffusive behavior in non-uniform crowded milieu by accounting for the convective transport of the macromolecules in crowded media. By this equation, the enhanced-diffusive transport under gradient is connected to the hindered diffusive transport which is well-reported and established in literature. Additionally, it introduces a correction factor to the general Fickian diffusion relation in cases of crowded environment to incorporate hard-sphere repulsive interaction between the species. For very small molecules, i.e. point particles, or at diluted concentrations, hard-sphere interaction is negligible and hence *D*_*eff*_ = *D*_0_, which is also predicted by equation 6.

We have experimentally observed tracer dependent transports in two different cases of crowding. The macromolecules have shown hindered-diffusion in uniformly-crowded environment. However, their transport was increased in non-uniformly-crowded medium, following an enhanced-diffusive pattern of transport, due to the repulsive interaction arising out of entropic volume exclusion driven hard sphere repulsions. We could successfully link both the diffusion types through a master relation, equation 6. Equivalent to the hydraulic buoyancy acting on a submerged object in a fluid, by reformatting equation 2, we can define diffusiophoretic buoyancy force as a driving force acting on any colloidal particles to push them out of regions of high “crowding pressure”. (see SI)

## Discussion

We have measured the transport of tracers in a gradient of crowder molecules by using a well-controlled microfluidic system. In contrast to the common techniques for studying diffusion under uniform crowder concentration, we explored the effect of the hard-sphere repulsive interaction on the transient dynamics of colloidal and molecular movements under crowder gradients which mimic biological media more accurately. We surprisingly observed that the transport of hard and soft tracers enhances in non-uniform crowded environments while dampens in uniform crowder concentrations. We further looked at this transport enhancement by varying the size and concentration of crowding species, as well as the size and type of tracers. We found that soft tracers experience a greater transport enhancement compared to hard tracers because the soft macromolecules and proteins can deform and squeeze through the interstitial spaces of the crowder. We hypothesize that hard-sphere repulsive interactions at higher crowder concentration cause the enhanced transport, in agreement with a hard-sphere diffusiophoretic model. This gives an effective “diffusiophoretic buoyancy force” which leads to the tracers escaping from the region with higher concentration of crowders (equation 2). The experimental data are consistent with this hypothesis for tracers both larger and smaller than the crowder.

Overall, our experiments and model provide a new perspective on transport in non-uniformly crowded environments that is applicable to various non-equilibrium systems, including biological environments such as the cell cytoplasm, where macromolecular crowding gradients are ubiquitous. Counterintuitively, we find *higher* transport rates of tracer particles in a gradient of crowder molecules. Thus, the intracellular molecular transport in living systems is not necessarily dampened in crowded environments as is commonly assumed. This new finding can lead to future theoretical work towards modelling more complicated, heterogeneous media such as protocells or synthetic cells which have multiple types of crowders. Also, this prompts us to study transport in “out-of-equilibrium” active crowded systems. We speculate that the observed effect, enhanced transport of colloids in crowded environments, is even higher in the biofluid of living species due to the higher mobility of active species^37,38^. The cytoplasm in a living cell is a broth of active constituents such as enzymes, metabolons and organelles that mix and agitate the surrounding fluids. The resulting hydrodynamic flows not only generate self-propulsion known in active systems, but can also increase the transport of other passive species. Interestingly, Takatori et al. proposed that active species generate an additional pressure, known as “swim pressure”, due to their activity ^39^. So, we expect that active crowders in biofluids would have increased “crowding pressure” compared to a solution of passive crowders, making the observed phenomenon especially relevant to living systems.^37,38^

## Acknowledgments

The work was supported by Penn State MRSEC funded by the National Science Foundation (DMR-1420620) and by NSF CBET-1603716.

## Author Contributions

DV and AS conceived the project. PB and THL guided the FCS experiments. MC, FM, SG carried the experiments. FM and MC did the modeling. All authors contributed to the writing of the manuscript.

## Additional Information

The authors declare no competing interests.

## Methods

### Fluorescence correlation spectroscopy (FCS) experiments

A custom-built microscope based optical setup was used to record the FCS signals. The excitation source in the microscope consisted of 532 nm, 80 MHz, 5.4 ps pulsed laser (High-Q Laser) from PicoTRAIN. The green laser is aligned to pass through an IX-71 microscope (Olympus), which was equipped with an Olympus 60× 1.2-NA water-immersion objective. Fluorescence signal from the samples were passed through a dichroic beam splitter (Z520RDC-SP-POL, Chroma Technology) and then focused onto a 50 μm, 0.22-NA optical fiber (Thorlabs), which acted as a confocal pinhole. The signal detected by the photomultiplier tube was routed to a preamplifier (HFAC-26) and subsequently fed to a time-correlated single-photon counting (TCSPC) board (SPC-630, Becker and Hickl). The setup on which the sample was placed is aligned with a high-resolution 3-D piezoelectric stage (NanoView, Mad City Laboratories)^40,41^.

The samples were excited using a constant and optimum laser power of ~30 ± 0.5 μW in order to prevent their photo-bleaching. 0.2 μM Rhodamine B solution was used to align the confocal pinhole and 50 nm fluorescent polystyrene particles (Fluoro-Max Fluorescent Polymer Microspheres, 1% solid, Thermo Fisher Scientific) was prepared in de-ionized water as reference standard (D = 9.5 × 10^−8^ cm^2^/s in water) for calibration^41^. The fluctuations in the sample signal arising from the in and out movement of the molecules with respect to the observation volume, were collected in first-in, first-out (FIFO) mode by the TCSPC board and fed into the instrument for detection. These fluorescence fluctuations were autocorrelated and employing Burst Analyzer 2.0 software, the data were fitted to a 3 dimensional model for obtaining the diffusion values. The corresponding values of τ_D_ were evaluated using the equation (1)^42,43^.

The Gaussian-shaped 3 dimensional observation volume with radial (r) and axial length (l) is used to obtain the normalized auto-correlation function, G(τ)^42,43^

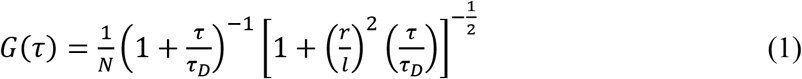

where the average number of particles in the observation volume is represented by N, τ denotes the autocorrelation time, τD is the diffusion time, and r/l indicates the structure factor.

The diffusion co-efficient of the 50 nm polystyrene reference standard at a temperature T was obtained from the Stokes-Einstein equation (2) and subsequently the confocal radius r was evaluated from the equation (3),^43^

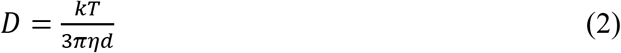

where D is the diffusion co-efficient, k is Boltzmann’s constant, η is the solvent viscosity and d is the size of the fluorescent samples.

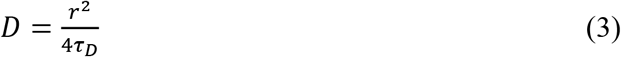

By evaluating the confocal radius r for the setup and the τ_D_ values from equation (1), the diffusion co-efficient of the samples can be precisely determined by equation (3)^43^.

For the FCS analysis, the initial step requires the dilution of the probe solution to nanomolar concentrations to ensure the existence of a single molecule in the observation volume. For studying the diffusion of fluorescently labelled particles in a crowded environment of the polymer, the particles were first diluted in 1 mM KCl solution, subsequently added to the polymer solutions and mixed thoroughly. The mixture was then subjected to sonication for ~15 minutes and this process was repeated before each sample preparation. 50 μL droplet of the sonicated sample solution was placed on a cover slip and FCS measurement for each samples were repeated 5 times. Finally, the diffusion co-efficient was obtained from the autocorrelation curves of the samples by fitting the curves with the one-component diffusion model.

### Microfluidic Channel Fabrication

We used soft lithography to fabricate the polydimethylsiloxane (PDMS) microfluidic channels in these experiments. Photolithography and etching were used to create the microchannel masters on silicon wafers in the Nanofabrication Laboratory of Materials Research Institute at the Pennsylvania State University. A M4L plasma etch was used to clean the wafers at 350 watts for 1 minute with 200 sccm of O2 and 50 sccm of He. Hexamethyldisilazane (HMDS) was spin coated on the wafers as an adhesion layer at 500 rpm for 15 seconds, 2500 rpm for 15 seconds and baked at 100°C for 30 seconds. Afterwards, we spin coated 5 mL of SPR 955 (Megaposit) onto the wafers at 500 rpm for 10 seconds, 750 rpm for 1 second, and 1500 rpm for 35 seconds, and baked at 100°C for 2 minutes. A second layer of SPR 955 was added to the wafer by repeating the previous step but was instead baked at 100°C for 1 minute. During photolithography, the mask with the microfluidic geometry was set on top of the photoresist covered wafer with a Karl Suss MA/BA6 Contact Aligner to expose the resist to UV radiation for 2 cycles of 8 seconds followed by baking at 100°C for 1 minute. Note that the microfluidic geometry on the mask was modeled in CAD and printed on a chrome-on-glass mask (Nanofabrication Laboratory, Materials Research Institute, Pennsylvania State University). Unexposed SPR 955 was removed by placing the wafers in MF CD26 developer for about 2 minutes, followed by washing with deionized water. The wafers were dried with nitrogen gas and then exposed to deep reactive ion etching (Alcatel) until the microchannel masters had a depth of 50 μm. Nanoremover PG (Micro Chem) was used to strip any remaining resist. The microchannel masters were silanized with Sigmacote (Sigma Aldrich) to prevent any adhesion when removing the microfluidic channel from the master pattern. PDMS (SylgardTM 184, Dow Corning) elastomer solution was prepared by combining the prepolymer and cross-linking agent in a weight ratio of 10:1 and poured over the microchannel masters to the desired thickness. The PDMS mixture was degassed in a vacuum desiccator for 1.5 hours to remove any air bubbles and then cured in an oven at 60 °C for 3 h. Next, the PDMS devices were removed from the microchannel masters and the inlets/outlet were punched out with a micro drill press. The resulting channels were adhered to glass slides (VWR) by exposing both the channels and glass slides to oxygen plasma and manually pressing them together (PDMS channel on top of the glass slide). After 24 hours to ensure proper sealing, polyethylene tubes (Becton Dickinson and Company, internal diameter of 0.61 mm) were connected to the drilled inlets/outlet to introduce fluid flow though the microchannels.

### Analysis of the Confocal Videos for Microfluidic Experiments

For the experiments of transport in crowder gradients in a microfluidic channel, 5-minute videos (633 frames, 475 ms/frame) were taken with confocal microscopy at the end of the channel at around 39 mm (17.4 s of residence time) at the mid-depth (~20-25 μm) as described in the main manuscript. There were 3 of these videos taken for control (without crowder) as well as 3 videos taken for experiment (with crowder). The 633 frames from one video was averaged by a z stack in ImageJ and a cut line window of (330 μm X 0.1 mm) was used to obtain the tracer distribution across the channel. The analysis was repeated to yield 3 tracer profiles for the control case and 3 tracer profiles for the experiment case, which are used to calculate the enhancement in transport distance. Note that the tracer profiles shown in Figure 1A are the average profile of the 3 tracer profiles for control and 3 tracer profiles for experiment.

In order to calculate the enhancement in transport distance, the slope was calculated from 7 data points near 0.5 fluorescence intensity on the same side to yield 3 slopes for control and 3 slopes for experiment. The inverse of these slopes was taken and then averaged to yield values for the lateral transport of 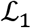 (control) and 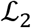 (experiment). The reported error is based on the 90% confidence interval calculated with the sample standard deviation of the three values for each 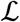. The enhancement in transport distance was calculated with 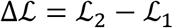 in μm.

### Modelling

Modelling has been done numerically in COMSOL Multiphysics using both modules of “Laminar Flow” and “Transport of Diluted Species”. To obtain the fluid flow in the channel, we initially solved the incompressible Navier-Stokes equation over the 3-domentional domain of the channel geometry (similar to figure 1A). No-slip and constant flow rate, 50*μl*/*hr*, were used for the boundary conditions at the channel walls and the three inlets, respectively. For highest accuracy, the effect of the viscosity has been taken into account by incorporating the experimentally-measured viscosities of the middle and side-streams. Viscosity of the dextran solution at different concentrations was measured using a viscometer (Brookfield DV-II+ Pro) (see SI for the viscosity of dextran 1000kDa). After obtaining the velocity profile of the fluid flow inside the channel, **u**_**h**_, the mass transport equation is solved under steady-state condition on the 3-D domain to obtain the polymer concentration profile, c:

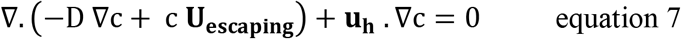

Where *D* is polymer diffusion as a function of concentration (equation 3), ∇ is the gradient operator, **U**_**escaping**_ is the escaping velocity defined in equation 2, and **u**_**h**_ is the fluid flow profile in the channel. After solving the equations for control experiment, 2wt% and 4wt% dextran, the concentration profile at the end of the channel were plotted together with the experimental result in Figure 6A.

