## Supplementary Information for "Non-uniform Crowding Enhances Transport: Relevance to Biological Environments"

### Contents

1. Mass Balance .......... S2
2. Initial Hydrodynamic Effects Upon the Addition of Polymer .......... S4
3. Relation Between Transport Distance and Absolute Diffusion .......... S6
4. Effect of Different Tracer Charges .......... S7
5. Crowder in the Middle Flow vs. in the Side Flows .......... S8
6. Initial uniform concentration of tracers with non-uniform crowder .......... S10
7. Size and Diffusion Data from DLS and FCS .......... S11
8. Interstitial size in crowder solutions .......... S12
9. Change of Radius of Gyration in crowder .......... S13
10. Diffusiophoretic Buoyancy .......... S14
11. Dextran Diffusion measurement using microfluidics .......... S15
12. Viscosity of dextran 1000kDa .......... S16
13. References .......... S17

### 1. Mass Balance

In this section we want to show that the conservation of mass holds for the corresponding non-normalized curve of Figure 1B, which is the normalized intensity profile of 60nm cPSL particles with and without PEO 1000kDa. Figure S1 is the corresponding intensity curve before normalization.

The mass balance equation for the tracer species can be written as the following:

$$\dot{m} = \int_0^W c.v.H\ dy \qquad \text{Equation S1}$$

where $c$ is concentration of tracer, $v$ is the velocity of the fluid along the channel, H is the height and W is the width of the channel. In the control case, the fluid velocity is relatively constant across the channel which can be calculated to be $v = 2.3\ \text{mm/s}$ based on the overall flow rate from the three inlets, $3 \times 50\ \mu\text{l/hr}$, and the cross-sectional area of the channel, $50 \times 360\ \mu\text{m}$. Upon addition of the polymer into the solution in the middle inlet, the viscosity increases which leads to the hydrodynamic spreading of the mid-solution from $\sim 120$ to $\sim 200\ \mu\text{m}$ (figure S3). Under this condition, each side flow has only $\frac{360-200}{2} = 80\ \mu\text{m}$ of the channel width to pass through, despite the similar flow rate of $50\ \mu\text{l/hr}$. This would force the side flows to have a faster velocity which can be calculated to be $\sim 3.45\ \text{mm/s}$. The middle flow also slows down due to a higher available width to pass through, having an average velocity of $\sim 1.4\ \text{mm/s}$. Having these average velocities and assuming that the fluorescent intensity is linear with concentration [1], we can calculate the result of the above integral for both of the curves shown in Figure S1. The result is shown in the table below. Note that before calculating the integral, the background intensity, the minimum intensity that the confocal still receives even when there is no fluorescent species present, needs to be subtracted from the intensity curves. As can be seen, the conservation of mass holds very well with $< 1\%$ error.

**Table S1: Calculated values of the conservation of mass integral (equation S1) for two cases shown in Figure S1C.** The very small difference shows that the conservation of mass holds very well with $< 1\%$ error.

| | Control Curve | Experiment Curve |
|---|---|---|
| **Minimum Intensity** | 8.5 | 8.2 |
| **Maximum Intensity** | 117.0 | 112.8 |
| **Integral** | 15343.2 | 15275.6 |
| **Difference** | 0.4% | |

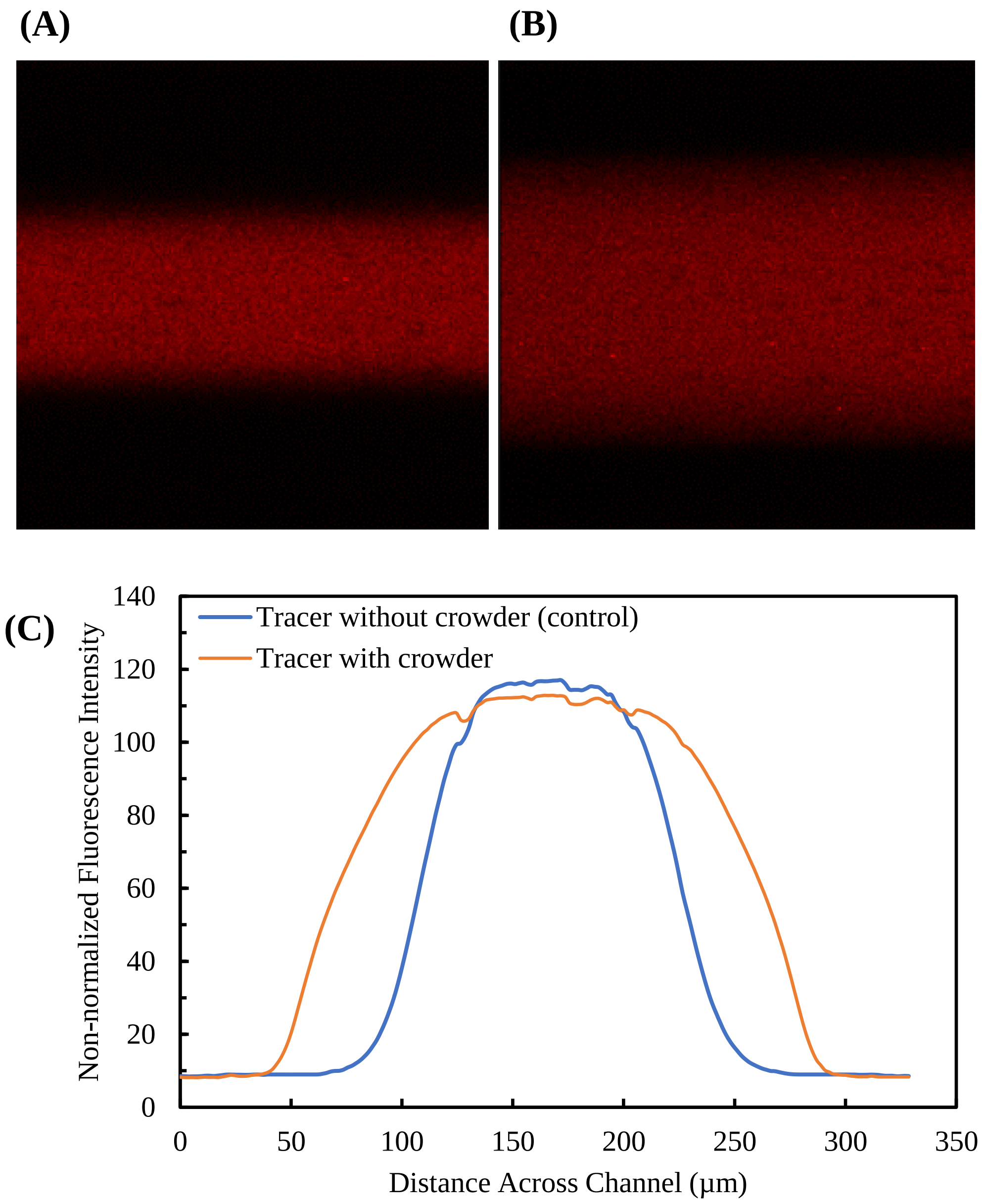


**Figure S1: Intensity profiles of 60 nm cPSL tracers at the end of the channel in the absence and presence of PEO 1000kDa crowder in the mid-inlet.** (A) Actual confocal image at the end of the channel showing the 60 nm cPSL tracers in the absence of crowder in the mid-inlet. (B) Actual confocal image at the end of the channel showing the 60 nm cPSL tracers in the presence of crowder in the mid-inlet. (C) The non-normalized intensity profiles of 60 nm cPSL tracers at the end of the channel in the absence and presence of PEO 1000 kDa crowder in the mid-inlet from (A) and (B).

### 2. Initial Hydrodynamic Effects Upon the Addition of Polymer

When the crowder is added to the microfluidic channel, there is an artificial broadening of the polymer phase as the polymer exhibits higher pressure due to having increased viscosity. Figure S2 shows the tracer particle profile taken at the beginning of the channel at around 0.5 mm right after mixing. One frame of the actual images from the beginning of the channel without (A) and with (B) crowder can be seen in Figure S3. In the tracer profile with crowder, the tracers have not had any time to interact with the crowder so that the tracer particles are effectively showing the crowder polymer phase. In both Figure S2 and Figure S3, the case without any crowder shows that the tracer phase is about 120 μm wide while the case with crowder shows that the crowder polymer phase is about 200 μm wide. The broadening of the polymer phase is due to the polymer exhibiting higher pressure due to having increased viscosity and not from tracer-crowder interaction. Note that the $\Delta\mathcal{L}$ at the beginning of the channel is 1.14 ± 1.15 μm or effectively 0 μm showing that the crowder has not had any time to repel the tracers and cause an enhancement in transport yet.

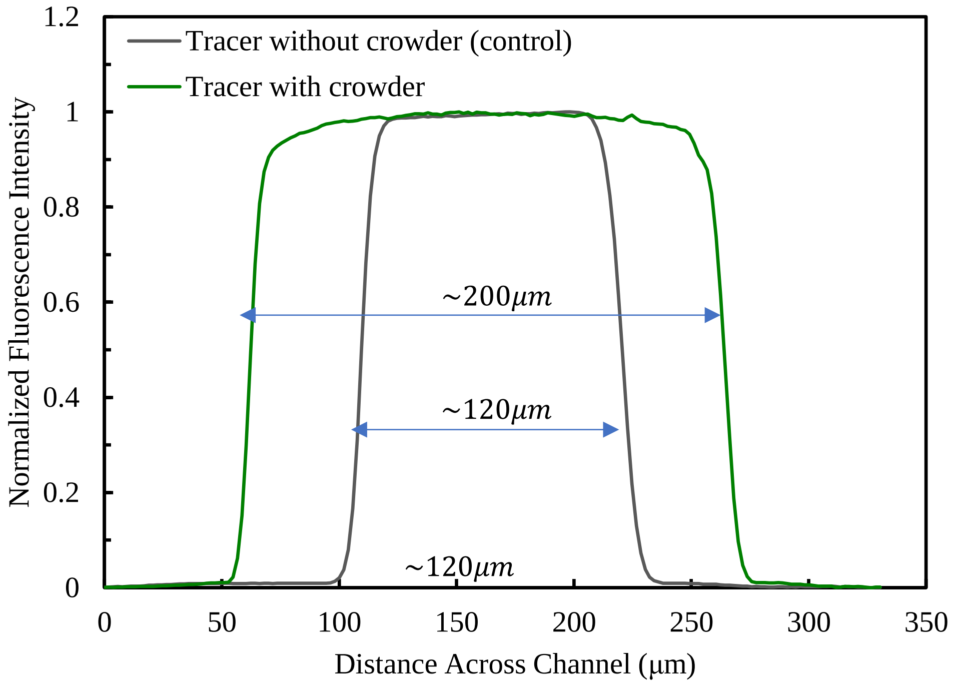


**Figure S2: The 60 nm cPSL tracer profile taken at the beginning of the channel without and with 0.3% 1000 kDa PEO crowder.**

(A)

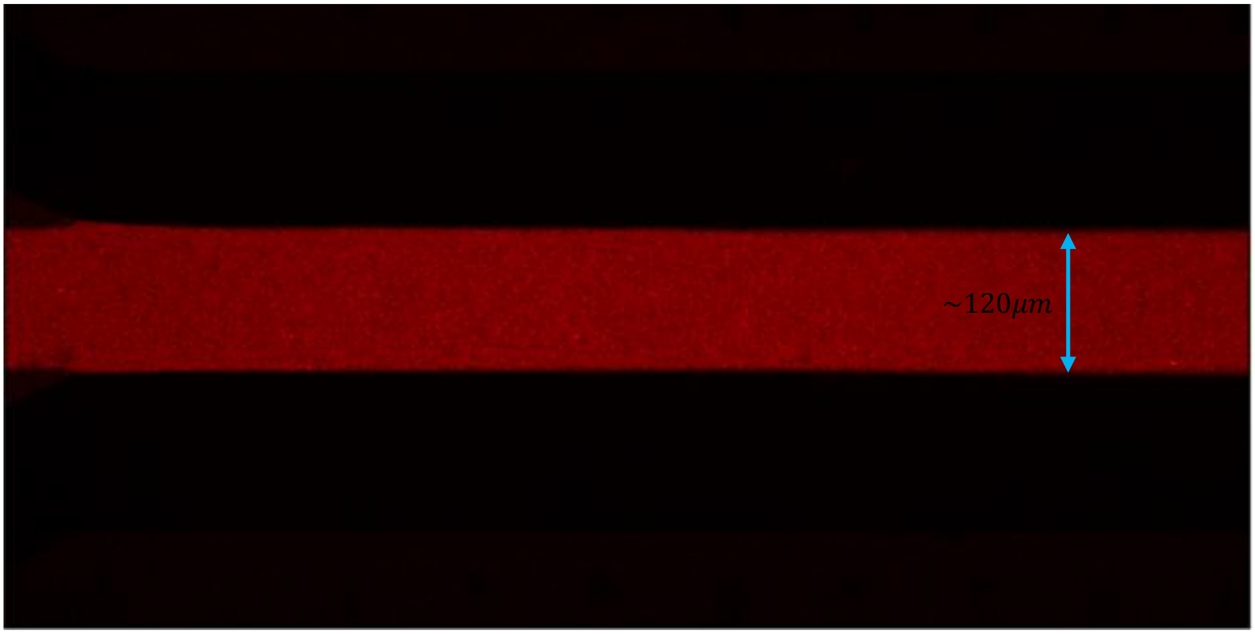


(B)

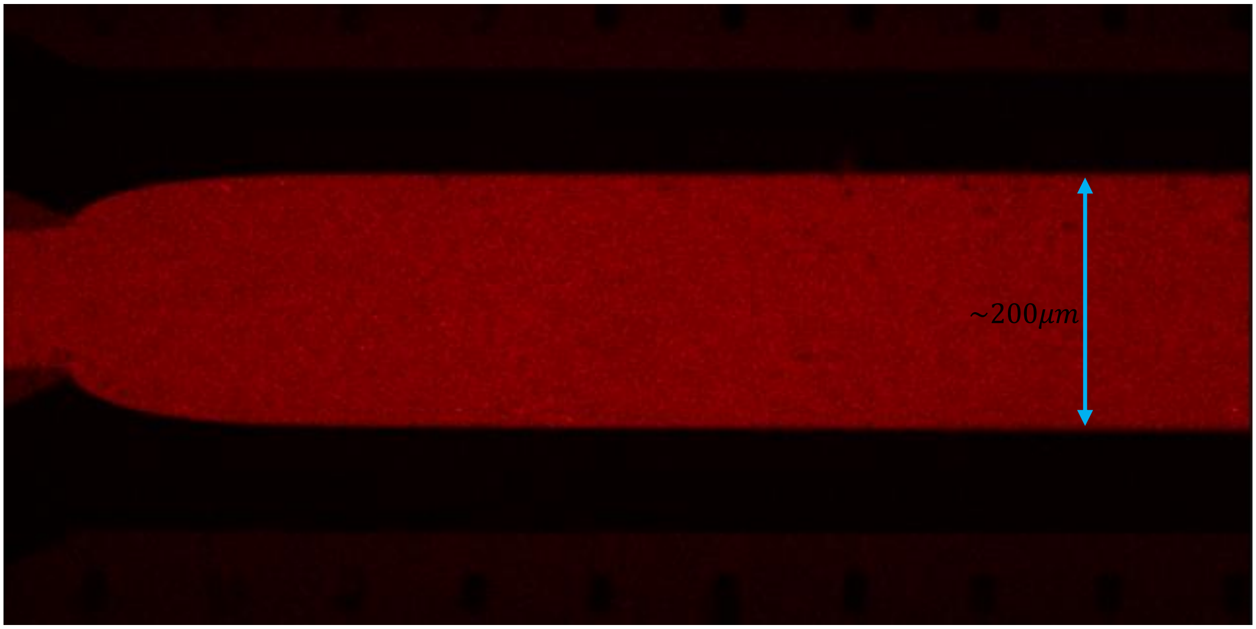


**Figure S3: Confocal images at the beginning of the channel showing the 60 nm cPSL tracers in the absence and prescence of crowder in the mid-inlet.** (A) absence of crowder in the mid-inlet (B) presence of crowder in the mid-inlet

### 3. Relation Between Transport Distance and Absolute Diffusion

In this section, the derivation of equation 1 in the manuscript is explained. This equation links the transport distance, $\mathcal{L}$, to the diffusion of the species in the channel. Several studies have shown that diffusion can be measured using T or H shaped microfluidics [2–5]. In a microfluidic channel, the flow is laminar due to a low Reynolds number, making diffusion the dominant mechanism of mixing between the parallel flows, i.e. across the channel. Along the channel however, the Peclet number is large and the diffusion is negligible. From scaling analysis, it is clear that the relation between transport distance, diffusion and residence time, $\tau$, should have the following form: $D \sim \mathcal{L}^2/\tau$. In our system, $\tau = 17.4$ secs. Using numerical and analytical calculation, we can determine the coefficient of this proportionality to be $\cong 1/4\pi$ [4,6,7]. Below we explain the numerical way.

When the tracer species is flown in the middle inlet, the tracers start to diffuse toward the sides as they travel along the channel (figure 1A). The height of the channel is much larger than the width of it, therefore we can simplify the 3D diffusion problem to 2D, along the length and width of the channel, without losing <0.1% accuracy [4]. Also, along the channel, $Pe_L = \frac{\bar{u}\ .\ L_{channel}}{D_{15nm\_silica}} \approx 10^6$ for the fastest diffusing species, meaning that we can easily neglect diffusion along the channel. $\bar{u}$ is the average velocity of the fluid inside the channel and $L_{channel}$ is the width of it. For more information, Häusler et al. have fully solved the full and simplified form of the diffusive-convective transport in microchannels and showed that the simplifications used here does not affect accuracy [4]. Under these simplifications, the equation for the transport of tracers becomes:

$$\bar{u}\frac{\partial C_{tr}}{\partial x} = D_{tr}\frac{\partial^2 C_{tr}}{\partial y^2} \qquad \text{Equation S2}$$

Where $C_{tr}$ and $D_{tr}$ are the concentration and diffusion coefficient of the tracers. $x\ and\ y$ is the coordinate across and along the channel, accordingly (figure 1A) [4]. Solving this equation numerically with the appropriate boundary conditions would give the concentration profile of the tracers, $C(x, y)$ from which $C(x = 39\text{ mm}, y)$ can be obtained. By taking the derivative, the slope of the concentration gradient can be found. Transport distance, $\mathcal{L}$, is defined as the inverse of the slope at the middle and side flow interfaces. By numerical simulation, we found that, as long as $\frac{\tau\ .\ D_{tr}}{L^2_{channel}} < 0.005$, $\mathcal{L}^2/D\tau \cong 4\pi$ which yields equation 1 in the main manuscript.

For this specific case shown in Figure S4, 60 nm cPSL tracers diffusing from the middle to the sides, the lateral transport distance is $\mathcal{L}_1 = 44.4 \pm 1.4$ µm which would give the diffusion coefficient of 9.0 $\pm$ 0.6 µm$^2$/s. This diffusion coefficient from microfluidics is close to the that obtained by FCS, 8.2 $\pm$

$0.5\ \mu m^2/s$ corresponding to $60 \pm 4$ nm in diameter. The size is consistent with the DLS measurement provided in Table S3 for cPSL.

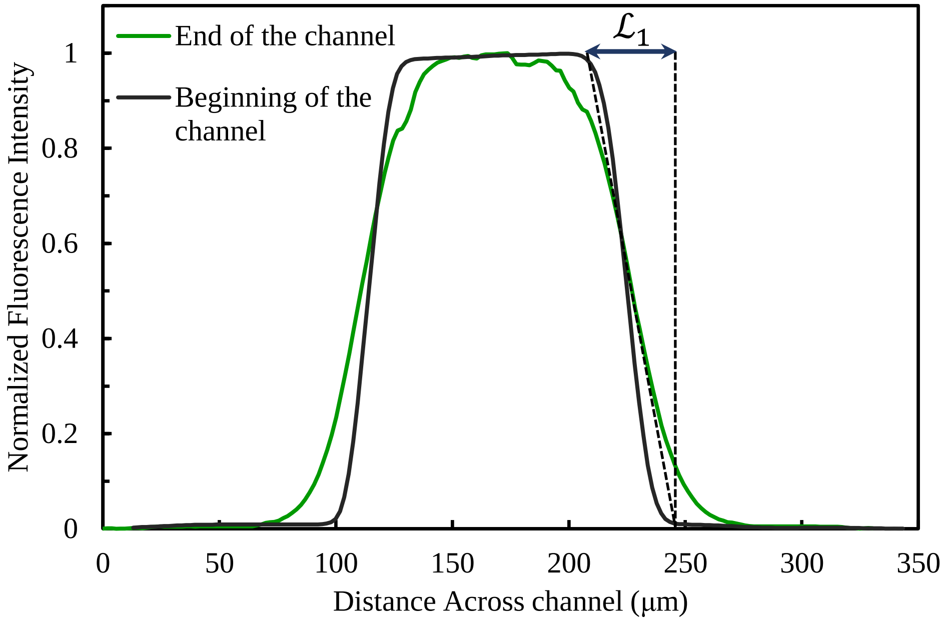


**Figure S4: Shows the distance that tracers diffused, $\mathcal{L}_1$, from the beginning to the end of the channel.** The tracer diffusion coefficient can be calculated based on this distance using equation 1.

### 4. Effect of Different Tracer Charges

To further investigate the effect of charge on the observed enhancement in the transport of tracers, we tested particles with different charges but similar sizes. We compared the $\Delta\mathcal{L}$ of similarly sized polystyrene spheres that were positively (aPSL, ~81.7 nm) and negatively charged (sPSL, 89.7 nm) in Figure S1. We saw that both particles show a $\Delta\mathcal{L}$ of around 14 μm, which shows that the particle charge does not influence the hard sphere interaction with the PEO crowder.

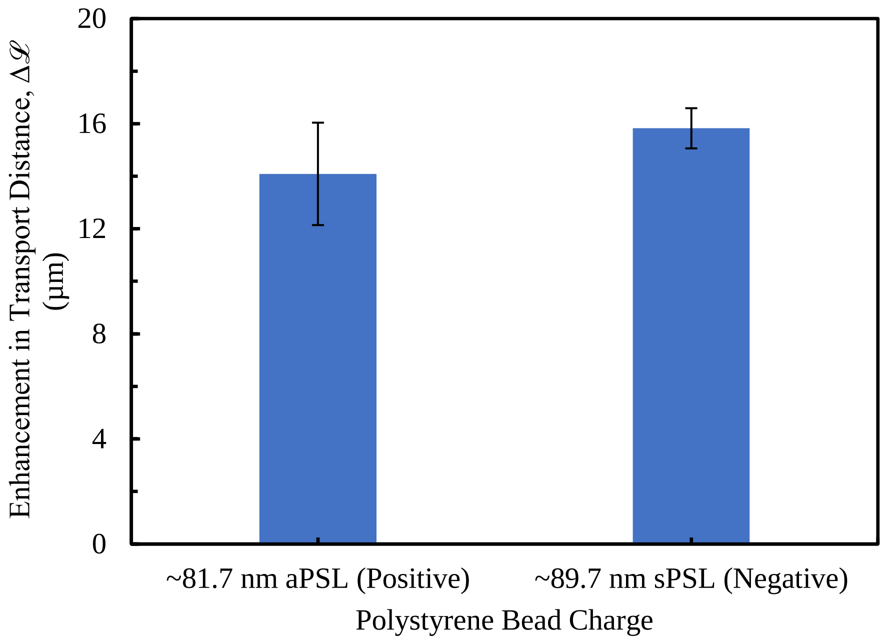


**Figure S5: Changing the charge of the polystyrene tracer particle does not change the observed enhancement in transport distance.**

### 5. Crowder in the Middle Flow vs. in the Side Flows

Here we show the effect on tracer transport when the crowder is introduced in different locations in the microfluidic channel. Specifically, Figure S5A shows the tracer particle profile when the crowder is introduced on the sides, while Figure S5B shows the enhancement in tracer transport comparison of crowder in the middle flow vs crowder in the side flows. In the case of crowder in the side flows, the tracers are repelled closer to the center of the channel resulting in a larger slope, while the tracers are repelled to the walls of the channel resulting in a smaller slope in the case of crowder in the middle flow. In both situations, the enhancement in transport is calculated by $\Delta\mathcal{L} = \mathcal{L}_2 - \mathcal{L}_1$ yielding the same absolute value of $\Delta\mathcal{L}$ with opposite signs. Therefore, the crowder causes similar effects on tracer transport no matter where the crowder is located.

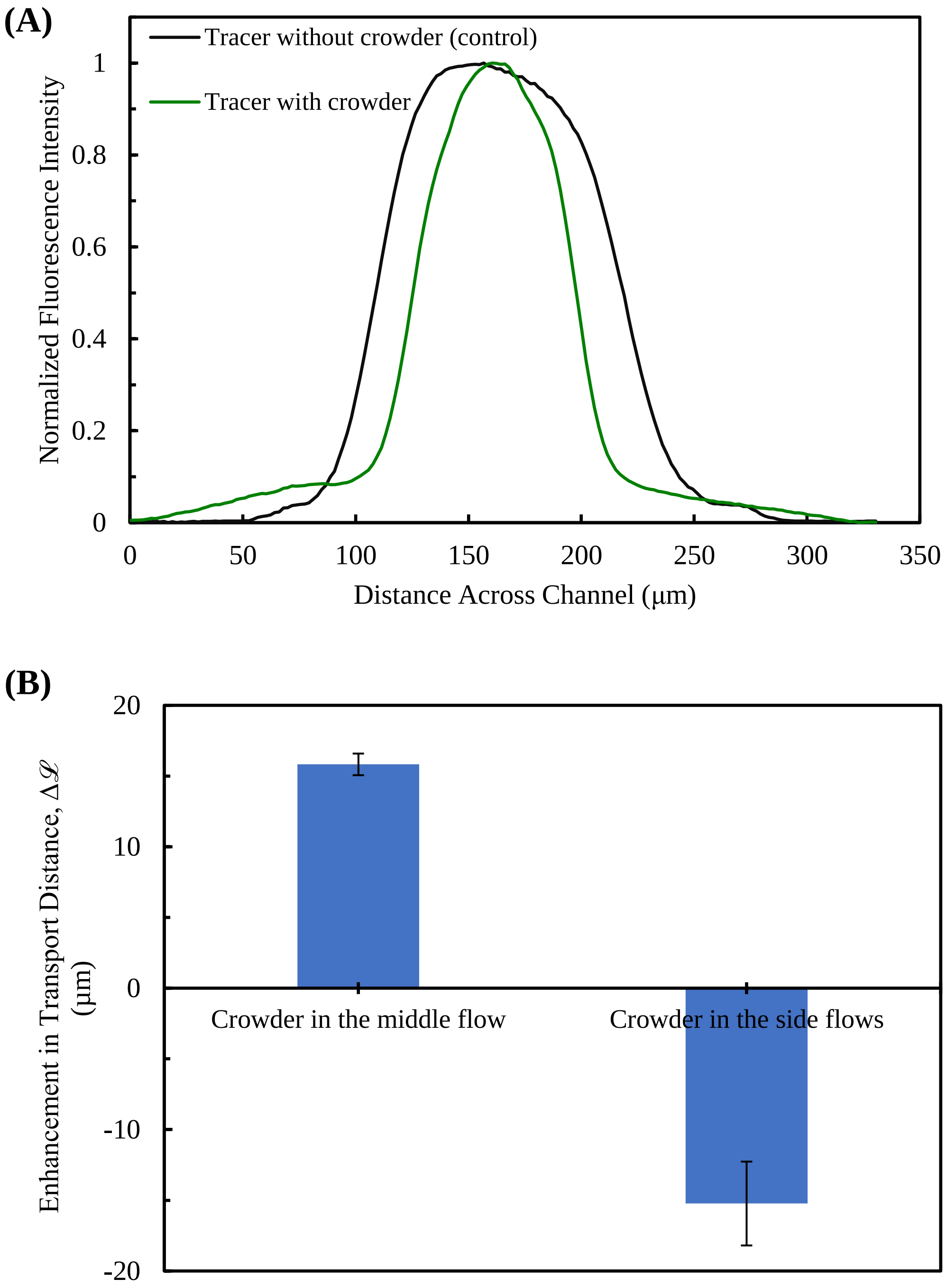


**Figure S6: The effect of crowder in the side flows on the tracer transport. (**A) The tracer particle profile when 0.3% PEO-1000kDa is in the side flows. There is no crowder in the middle flow. (B) The enhancement in transport distance, $\Delta\mathcal{L}$, for the cases where crowder originates in the middle flow and crowder originates in the side flows. Both cases show a $\Delta\mathcal{L}$ that is similar in magnitude but opposite in sign as the crowder is repelling the tracers in different directions.

### 6. Initial uniform concentration of tracers with non-uniform crowder

Here, we demonstrate the experimental data obtained from performing the microfluidic experiment (Figure 1A) for a specific case where we start with uniform concentration of tracers at the beginning of the channel (all three inlets) with crowders present only in the side inlets. The tracer is fluorescently tagged Dextran-2000 kDa at an infinitesimal concentration (0.02wt%) and the crowder is PEO-1000kDa at 0.3wt%. The normalized intensity profiles of the tracers are shown in Figure S7 at the beginning and the end of the microfluidic channel. Initially, the concentration of the tracers is equal everywhere (black curve), however, the profile is non-uniform near the end of the channel, with the highest concentration in the middle flow of the channel accompanied by two dips in concentration on either side. This intensity profile is a clear indication of tracer transport (focusing) from the region with high level of crowding (sides) to the dilute crowder location (middle flow).

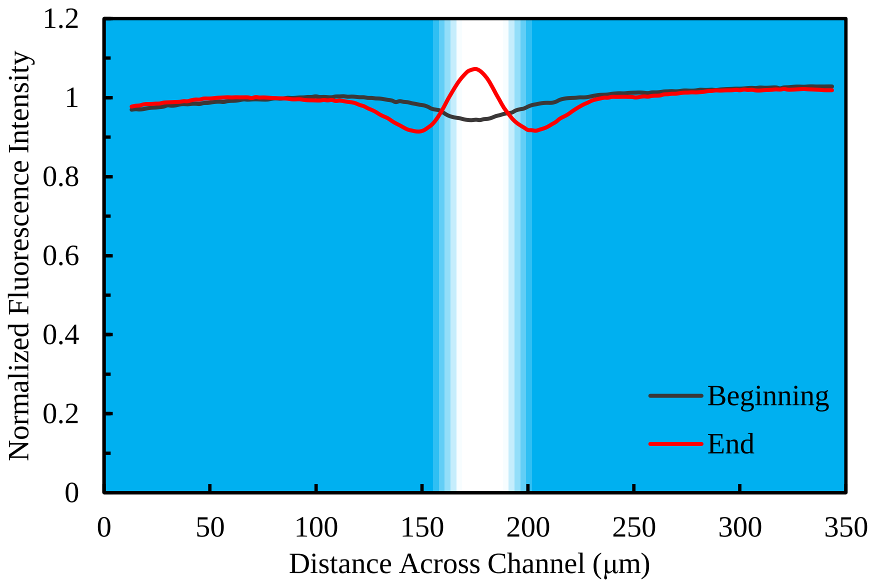


**Figure S7: Intensity curves of polymeric tracers at the beginning and the end of the microfluidic channel when the experiment is done under the specific condition: uniform initial concentration of tracers at the beginning with a non-uniform crowder distribution.** Dextran-2000kDa is the polymeric tracer present in all three inlets and PEG-1000kDa at 0.3wt% only in the side flows (blue background shade).

### 7. Size and Diffusion Data from DLS and FCS

We used PEG/PEO solutions of different molecular weights as crowders. Using DLS, we measured the average diameter size of the PEG/PEO molecules. Also, based on the molecular weight, nominal radius of gyration ($R_g$) and hydrodynamic radius ($R_h$) as well as overlapping concentration ($C^*$) are calculated for different PEG/PEO molecular weights using the below equations:

$$R_g(nm) = \frac{R_h}{0.65} = 0.0215\,MW^{0.58} \qquad \text{Equation S3}$$

$$C^*(gr/m^3) = \frac{MW}{4/3\,\pi\,R_g^3\,N_A} \qquad \text{Equation S4}$$

where MW is molecular weight, and $N_A$ is the Avogadro number. The $R_g$, $R_h$ and $C^*$ are calculated and provided in table S2. As can be seen, the measured DLS size is close to the nominal hydrodynamic diameter, $2R_h$, for each molecular weight PEG/PEO.

**Table S2: The sizes of the PEG/PEO crowding polymers from DLS measurements with the overlapping concentrations**

| PEG/PEO crowding species MW (kDa) | DLS diameter size (nm) | Calculated $R_g$ (nm) | Calculated $R_h$ (nm) | $C^*$ (wt %) |
|---|---|---|---|---|
| 20 | 12.7±1.6 | 6.7 | 4.4 | 2.62 |
| 200 | 43.7±5.3 | 25.5 | 16.6 | 0.48 |
| 600 | 68.9±3.2 | 48.3 | 31.4 | 0.21 |
| 1000 | 96.9±9.1 | 64.9 | 42.2 | 0.14 |
| 2000 | 116.0±8.9 | 97.1 | 63.1 | 0.09 |

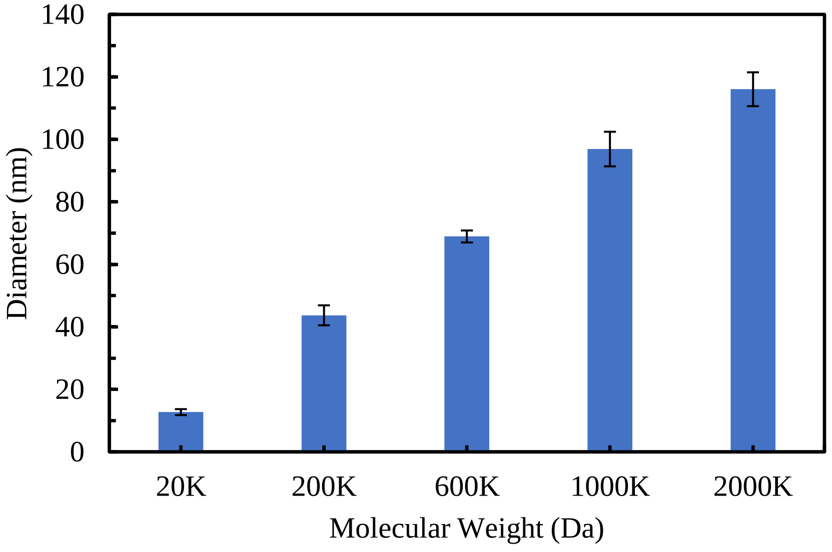

**Figure S8:** The sizes of the PEG/PEO crowding polymers from DLS measurements as given in Table S2

**Table S3: The sizes of the tracers from DLS and FCS measurements.** cPSL: carboxylate-modified polystyrene particles, aPSL: amine-modified polystyrene particles, sPSL: sulfate-modified polystyrene particles.

| Method | Tracer Species | Diffusion ($\mu m^2/s$) | Diameter size (nm) |
|---|---|---|---|
| FCS | 15nm Silica particles | $33.6 \pm 4.8$ | $14.6 \pm 2.1$ |
| | 20nm cPSL | $14.2 \pm 0.5$ | $34.4 \pm 1.3$ |
| | 60nm cPSL | $8.2 \pm 0.4$ | $59.6 \pm 2.9$ |
| | 100nm aPSL | $6.0 \pm 0.6$ | $81.7 \pm 8.8$ |
| | FITC Dextran 2000kDa | $7.6 \pm 0.6$ | $64.7 \pm 5.3$ |
| | Urease | $29.2 \pm 1.1$ | $16.8 \pm 0.7$ |
| | Aldolase* | $44.5 \pm 1.5$ | $11.0 \pm 0.4$ |
| DLS | 60nm cPSL | - | $61.0 \pm 0.4$ |
| | 100nm sPSL | - | $89.7 \pm 0.5$ |
| | 500nm sPSL | - | $276.2 \pm 3.0$ |
| | 1µm cPSL | - | $542.8 \pm 8.5$ |
| | FITC Dextran 500kDa | - | $39.1 \pm 1.2$ |
| | Vesicles (extruded) | - | $221.8 \pm 4.2$ |

### 8. Interstitial size in crowder solutions

In semidilute regime of polymer concentration, the solution space is filled with correlation blobs with correlation length $\xi$ as follows:

$$\xi = R_g^0 (C/C^*)^{-\nu/(3\nu-1)} \qquad \text{Equation S5}$$

Where, $R_g^0$ is the radius of gyration in dilute solution, $C$ and $C^*$ are the polymer (crowder) concentration and critical concentration, respectively and $\nu$ is the Flory exponent which is ~ 0.588 for a good solvent case.

We could calculate $\xi$ ~ 40 nm for 0.3% PEO1000 kDa crowder solution which was used in experiments shown in Figure 4. The average interstitial size which could be thought of as the characteristic size between correlations blobs, could be calculated based on 3D packing of such blobs or spheres.

Mathematically, the interstitial size would be ~ (√3-1)×40 ~ 29 nm. Experimentally, we also found that the transport enhancement dramatically increased for hard sphere tracers around this interstitial size. Below this length scale, hard sphere tracers cannot possibly feel the volume exclusion effect being able to fit in efficiently and soft tracers can squeeze in size to fit in more efficiently and thereby, increase the enhanced transport (Figure 4A). Overall, volume exclusion only works on hard tracers, if tracer size is comparable to the interstitial size.

### 9. Change of Radius of Gyration in crowder

Soft tracers show increased enhanced transport compared to hard tracers. This can be explained in the light of soft, squishy and deformable nature of soft tracers which are forced to decrease their hydrodynamic size due to volume fraction increase of the crowder polymer in the semidilute concentration regime.

In developing this model, we assumed (i) volume fraction effect is only coming from crowders and neglected the tracer volume fraction contribution and (ii) both the soft and the hard tracers feel the same viscosity (i.e. microviscosity given the same size of the tracer). Although this assumption partially holds and might deviate in cases of larger size tracers (like vesicles), we can get an estimate quite easily.

In semidilute regime of PEO crowder solution around the transition, the crowder volume fraction $\phi$ can be expressed as a function of its molecular weight MW as follows:

$$\phi = 900\, MW^{-0.76} \qquad \text{Equation S6}$$

The semidilute transition volume fraction $\phi^*$ can be expressed in terms of degree of polymerization, N, of the crowder:

$$\phi \sim N^{-4/5} \qquad \text{Equation S7}$$

The radius of gyration of a soft tracer (eg. dextran) in semi-dilute regime at volume fraction $\phi$ can be expressed as follows:

$$R_g = R_g^0(\phi/\phi^*)^{-0.125} \qquad \text{Equation S8}$$

The hydrodynamic radius ($R_h$) of soft tracer which describes self-diffusion coefficient of the tracer is related to the radius of gyration as follows:

$$R_h = 0.65\, Rg \qquad \text{Equation S9}$$

Note that for hard tracers, there is no statistical dimension or $Rg$ and $R_h$ is simply the radius of the hard sphere.

We estimate that for 0.3 wt% PEO1000 kDa crowder solution, the ratio of radius of gyration between soft and hard tracers, $R_g/R_g^0 \sim (\phi/\phi^*)^{-0.125} \sim (0.0248/(3.3 \times 10^{-4}))^{-0.125} \sim 0.58$.

Therefore, the ratio of diffusivity between soft tracer and hard tracer of the same size: $r_D \sim \frac{1}{0.65*0.58} \sim 2.66$.

Following this, the theoretical ratio of diffusive length scales between soft and hard tracers: $r_L \sim \sqrt{2.66} \sim 1.63$. Note that this ratio of diffusive length scale would also reflect into their ratio of transport enhancement length scales as shown in Figure 4A. From experimental results shown in Figure 4A, we could calculate the ratio of transport enhancement between soft and hard tracers as $r_L \sim \frac{25}{15} \sim 1.66$. Therefore, our theoretical model agrees with experimental data and observations.

### 10. Diffusiophoretic Buoyancy

According to the non-electrolyte diffusiophoresis for a colloidal particle, gradient of an interacting solute can induce a surface slip velocity which drags the particle toward or away from the high concentration region, depending on the attractive or repulsive nature of the interaction. The velocity of the particle can be calculated using the following equation:

$$\mathrm{U_{dp}} = \frac{\mathrm{kT}}{\eta} \mathrm{KL^*}\, \nabla \mathrm{C_{solute}} \qquad \text{Equation S10}$$

where $k$T is the thermal energy, $\eta$ is the viscosity of the solution; and $\nabla C_c$ is concentration gradient of the solute species. KL* is the parameter calculated based on the solute-colloid interaction, having positive value for attractive and negative value for repulsive interaction [8]. For hard-sphere repulsion, KL* is equal to $-r_c^2/2$ where $r_c$ is the radius of the solute crowder. Also, osmotic pressure in a region with crowder concentration of $C_c$ can be defined as $\Pi_c = kTC_c$. Therefore, the escape velocity of the colloidal particles from region of high crowder concentration can be obtained by:

$$\mathrm{U_{escape}} = -\frac{r_c^2}{2\,\eta} \nabla \Pi_c \qquad \text{Equation S11}$$

Which is similar to equation 2 in the manuscript. To stop the particle from moving, an opposing external force is required such that $F_{external} + F_{dp} = 0$. Due to the linearity of the Naiver-Stokes equation at low Reynold's number, the diffusiophoretic force can be obtained by:

$$F_{\mathrm{dp}} = -3\pi\, \mathrm{a}\, \mathrm{r}_c^{\,2}\ \nabla\Pi_{\mathrm{c}} \qquad \text{Equation S12}$$

Where a and $r_c$ are the radius of the tracer and crowder species. This form of equation is similar to the hydrostatic buoyancy force that acts on a submerged object with volume V and pushes it out of the region of high fluid pressure according to the following relation: $\mathrm{F_h} = -\mathrm{V}\,\nabla \mathrm{P_h}$. So, the diffusiophoretic force (eq. 12) has a similar form as hydrostatic buoyancy, it is the multiply of osmotic pressure gradient and an equivalent volume which we can refer to as effective diffusiophoretic volume:

$$V_{dp} = 3\pi\, \mathrm{a}\, \mathrm{r}_c^{\,2} \qquad \text{Equation S13}$$

### 11. Dextran Diffusion measurement using microfluidics

Here, the experimental data is shown for the case of measuring the hindered self-diffusion of fluorescent dextran in uniform concentration of crowder (untagged dextran-2000kDa at 2 and 4 wt%) inside a microfluidic channel. Figure S9A shows the normalized intensity of the tagged dextran in the absence of crowder (control) and in the uniform presence of 2 and 4 wt% of crowder everywhere. Figure S9B shows the obtained diffusion coefficient of the tracer based on the slope analysis of Figure S9A under the three crowding conditions. As can be seen, the diffusion data we obtained from microfluidic experiment is consistent with the measured value of diffusion using FCS (Figure 6B in the manuscript).

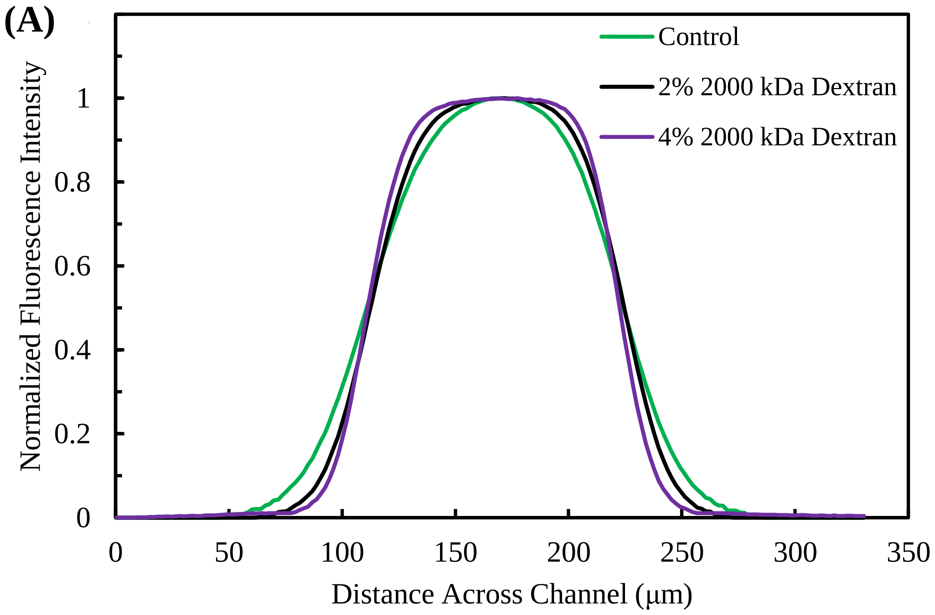

(B)

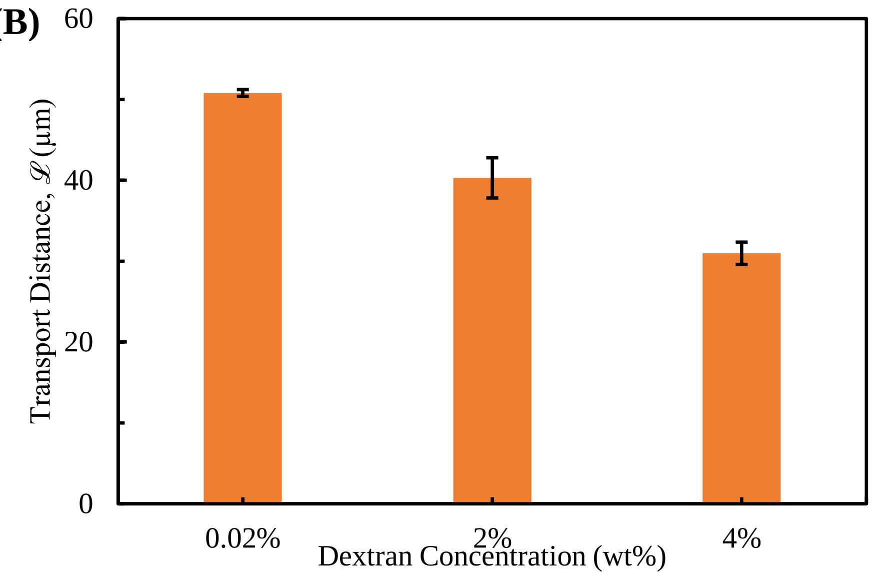


**Figure S9: The measurement of dextran diffusion using microfluidics where there is uniformly distributed crowder.** (A) The diffusion profile from the microfluidic channel of fluorescent 2000 kDa dextran in uniform 2 and 4 wt% untagged 2000 kDa dextran. The control is the diffusion profile of 0.02 wt% 2000 kDa in 1mM KCl buffer. (B) The transport distance, $\mathcal{L}$, of the diffusion profiles of the fluorescent 2000 kDa dextran from A.

### 12. Viscosity of dextran 1000kDa

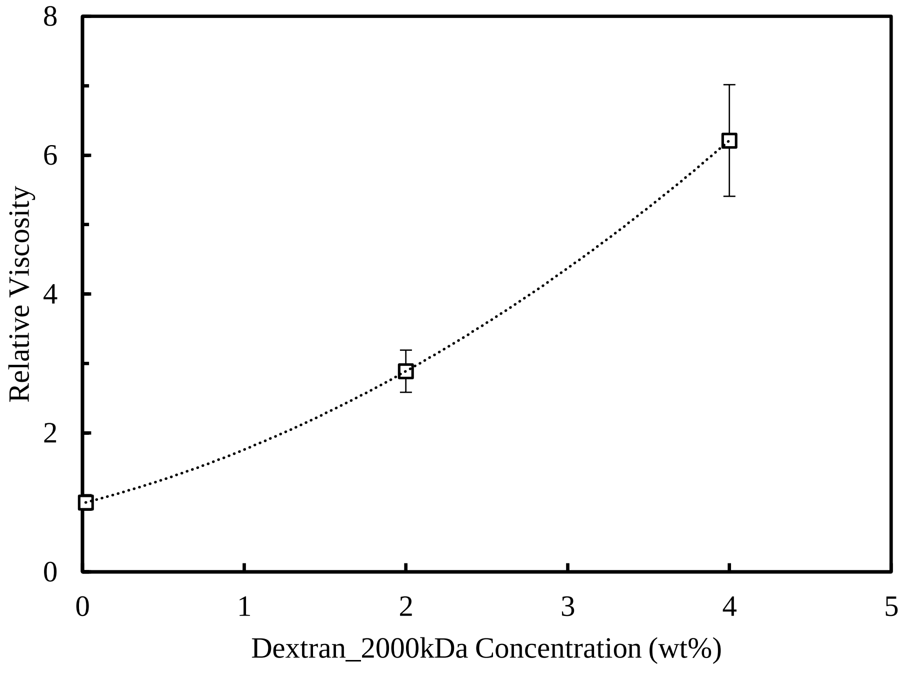


**Figure S9:** Relative viscosity of Dextran-2000kDa solution at different concentration.
